# A potential SARS-CoV-2 variant of interest (VOI) harboring mutation E484K in the Spike protein was identified within lineage B.1.1.33 circulating in Brazil

**DOI:** 10.1101/2021.03.12.434969

**Authors:** Paola Cristina Resende, Tiago Gräf, Anna Carolina Dias Paixão, Luciana Appolinario, Renata Serrano Lopes, Ana Carolina da Fonseca Mendonça, Alice Sampaio Barreto da Rocha, Fernando Couto Motta, Lidio Gonçalves Lima Neto, Ricardo Khouri, Camila Indiani de Oliveira, Pedro Santos-Muccillo, Joao Felipe Bezerra, Dalane Loudal Florentino Teixeira, Irina Riediger, Maria do Carmo Debur, Rodrigo Ribeiro-Rodrigues, Anderson Brandao Leite, Cliomar Alves do Santos, Tatiana Schäffer Gregianini, Sandra Bianchini Fernandes, André Felipe Leal Bernardes, Andrea Cony Cavalcanti, Fábio Miyajima, Claudio Sachhi, Tirza Mattos, Cristiano Fernandes da Costa, Edson Delatorre, Gabriel L Wallau, Felipe G Naveca, Gonzalo Bello, Marilda Mendonça Siqueira, on behalf of Fiocruz COVID-19 Genomic Surveillance Network

## Abstract

The SARS-CoV-2 epidemic in Brazil was dominated by two lineages designated as B.1.1.28 and B.1.1.33. Two SARS-CoV-2 variants harboring mutations at the receptor-binding domain of the Spike (S) protein, designated as lineages P.1 and P.2, evolved within lineage B.1.1.28 and are rapidly spreading in Brazil. Lineage P.1 is considered a Variant of Concern (VOC) because of the presence of multiple mutations in the S protein (including K417T, E484K, N501Y), while lineage P.2 only harbors mutation S:E484K and is considered a Variant of Interest (VOI). Here we report the identification of a new SARS-CoV-2 VOI within lineage B.1.1.33 that also harbors mutation S:E484K and was detected in Brazil between November 2020 and February 2021. This VOI displayed four non-synonymous lineage-defining mutations (NSP3:A1711V, NSP6:F36L, S:E484K, and NS7b:E33A) and was designated as lineage N.9. The VOI N.9 probably emerged in August 2020 and has spread across different Brazilian states from the Southeast, South, North and Northeast regions.

## Background

The SARS-CoV-2 epidemic in Brazil was mainly driven by lineages B.1.1.28 and B.1.1.33 that probably emerged in February 2020 and were the most prevalent variants in most country regions until October 2020 ^1,2^. Recent genomic studies, however, bring attention to the emergence of new SARS-CoV-2 variants in Brazil harboring mutations at the receptor-binding site (RBD) of the Spike (S) protein that might impact viral fitness and transmissibility.

So far, one variant of concern (VOC), designated as lineage P.1, and one variant of interest (VOI), designated as lineage P.2, have been identified in Brazil and both evolved from the lineage B.1.1.28. The VOC P.1, first described in January 2021 ^3^, displayed an unusual number of lineage-defining mutations in the S protein (L18F, T20N, P26S, D138Y, R190S, K417T, E484K, N501Y, H655Y, T1027I) and its emergence was associated with a second COVID-19 epidemic wave in the Amazonas state ^4,5^. The VOI P.2, first described in samples from October 2020 in the state of Rio de Janeiro, was distinguished by the presence of the S:E484K mutation in RBD and other four lineage-defining mutations outside the S protein ^6^. The P.2 lineage has been detected as the most prevalent variant in several states across the country in late 2020 and early 2021 (https://www.genomahcov.fiocruz.br).

Several B.1.1.33-derived lineages are currently defined by the Pangolin system including: lineage N.1 detected in the US, lineage N.2 detected in Suriname and France, lineage N.3 circulating in Argentina, and lineages N.4 and B.1.1.314 circulating in Chile. However, none of these B.1.1.33-derived lineages were characterized by mutations of concern in the S protein. Here, we define the lineage N.9 within B.1.1.33 diversity that harbors mutation E484K in the S protein as was detected in different Brazilian states between November, 2020 and February, 2021.

## The Study

Our genomic survey of SARS-CoV-2 positive samples sequenced by the Fiocruz COVID-19 Genomic Surveillance Network between 12th March 2020 and 27th January 2021, identified 422 sequences as belonging to the lineage B.1.1.33 (Appendix 1). Next, these sequences were combined with 816 B.1.1.33 Brazilian genomes available in the EpiCoV database in GISAID by 1^st^ March 2021 (Appendix Table 1). Mutational profile was investigated using the nextclade tool (https://clades.nextstrain.org), finding the S:E484K mutation in 34 sequences. Maximumlikelihood (ML) phylogenetic analysis conducted using IQ-TREE v2.1.2^7^ revealed that 32 B.1.1.33(E484K) sequences branched in a highly supported (approximate likelihood-ratio test [aLRT] = 98%) monophyletic clade that define a potential new VOI designated as N.9 PANGO lineage ^8^. The other two sequences harbouring the E484K mutation branched separately in a highly supported (aLRT = 100%) dyad (**Figure 1a**). The VOI N.9 is characterized by four non-synonymous lineage-defining mutations (NSP3:A1711V, NSP6:F36L, S:E484K and NSP7b:E33A) and also contains a group of three B.1.1.33 sequences from the Amazonas state that has no sequencing coverage in the position 484 of the S protein, but share the remaining N.9 lineage-defining mutations (**Table 1**). The B.1.1.33(E484K) dyad comprises two sequences from the Maranhao state and were characterized by a different set of non-synonymous mutations (**Data not shown**).

**Figure 1.**
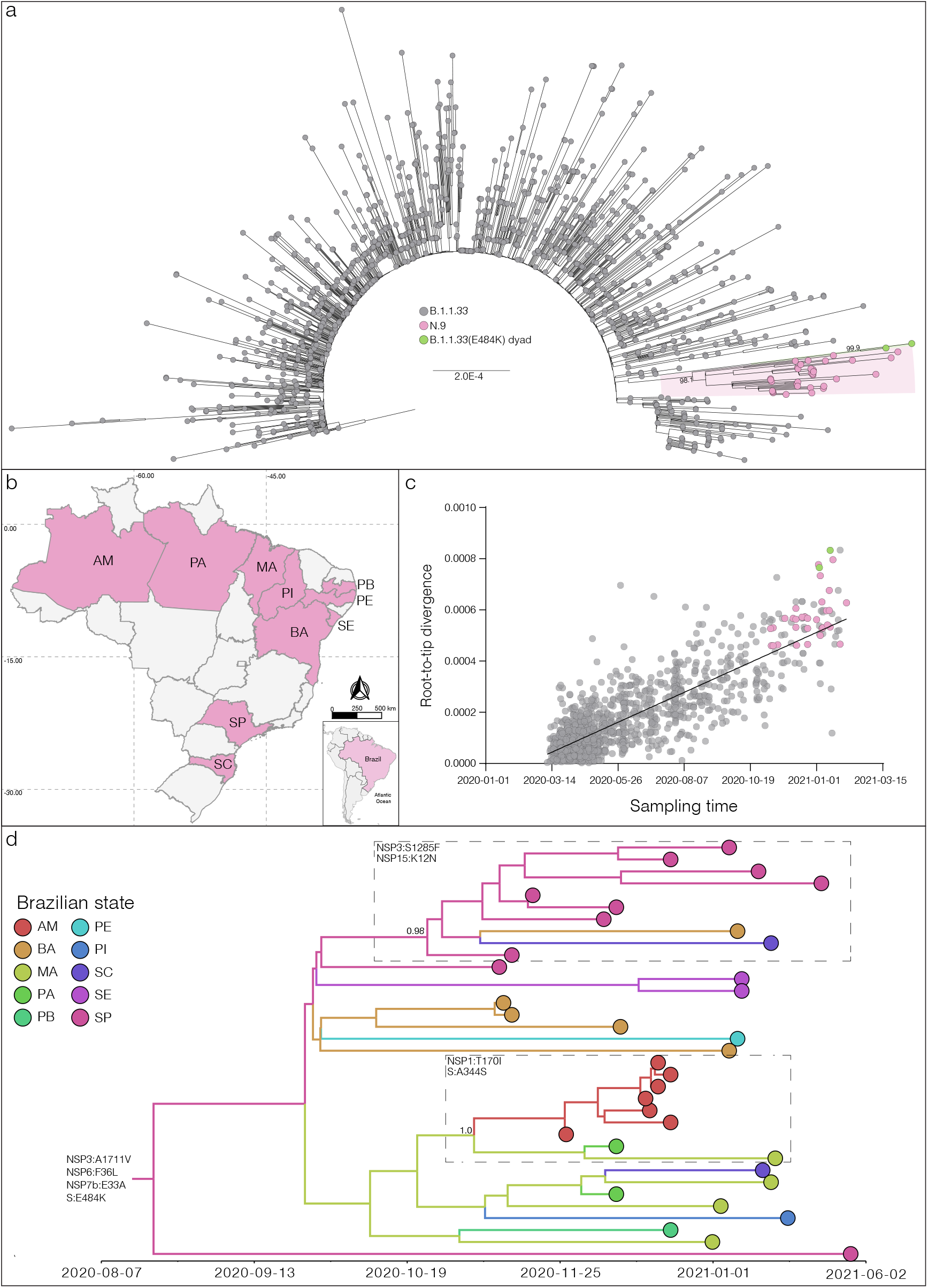
a) Maximum likelihood (ML) phylogenetic tree of the B.1.1.33 whole-genome sequences from Brazil. The B.1.1.33 sequences with mutation S:E484K are represented by pink (VOI N.9) and green [B.1.1.33(E484K)] circles. VOI N.9 clade is highlighted in pink. The aLRT support values are indicated in key nodes and branch lengths are drawn to scale with the left bar indicating nucleotide substitutions per site. b) Geographic distribution of the VOI N.9 identified in Brazil. Brazilian states’ names follow the ISO 3166-2 standard. c) Correlation between the sampling date of B.1.1.33 sequences and their genetic distance from the root of the ML phylogenetic tree. Colors indicate the B.1.1.33 clade as indicated in a). d) Bayesian phylogeographic analysis of N.9 lineage. Tips and branches colors indicate the sampling state and the most probable inferred location of their descendent nodes, respectively, as indicated in the legend. Branch posterior probabilities are indicated in key nodes. Boxes highlight two N.9 subclades carrying additional mutations (indicated in each box). The tree was automatically rooted under the assumption of a strict molecular clock, and all horizontal branch lengths are time-scaled.

**Table 1.** Synapomorphic mutations of SARS-CoV-2 lineage N.9.

| Genomic region (protein) | Nucleotide | Amino acid |
| --- | --- | --- |
| ORF1a | G1264T | - |
| ORF1a | C7600T | - |
| ORF1a (NSP3) | C7851T | A2529V (A1711V) |
| ORF1a (NSP6) | T11078C | F3605L (F36L) |
| Spike (S) | G23012A | E484K |
| ORF7b (NSP7b) | A27853C | E33A |

Among the 35 genomes identified so far as VOI N.9, 10 Brazilian states were represented, suggesting that this lineage is already highly dispersed in the country. The VOI N.9 was first detected in Sao Paulo state on November 11th, 2020, and soon later in other Brazilian states from the South (Santa Catarina), North (Amazonas and Para) and Northeast (Bahia, Maranhao, Paraiba, Pernambuco, Piaui, and Sergipe) regions (**Figure 1b**). Analysis of the temporal structure using TempEst ^9^ revealed that the overall divergence of lineage N.9 is consistent with the substitution pattern of other B.1.1.33 sequences (**Figure 1c**), thus suggesting no unusual accumulation of mutations in this VOI. Bayesian reconstruction using a strict molecular clock model with a uniform substitution rate prior (8 - 10 × 10^-4^ substitutions/site/year) as implemented in BEAST 1.10 ^10^ estimated the emergence of the VOI N.9 most probably in the states of Sao Paulo (*Posterior State Probability* [*PSP*] = 0.42), Bahia (*PSP* = 0.32) or Maranhao (*PSP* = 0.18) at 15^th^ August, 2020 (95% High Posterior Density [HPD]: 16^th^ June – 22^th^ September, 2020) (**Figure 1d**). This analysis also revealed that some additional mutations were acquired during evolution of VOI N.9 in Brazil, determining two highly supported (*PP* > 0.95) subclades. One subclade, that mostly contains sequences from Sao Paulo state, probably arose on 16^th^ October (95% HPD: 22^th^ September – 5^th^ November) and was defined by additional mutations NSP3:S1285F and NSP15:K12N. The other subclade that mostly comprises sequences from the North region probably arose on 29^th^ October (95% HPD: 5^th^ October – 17^h^ November) and was defined by additional mutations NSP1:T170I and S:A344S (**Figure 1d**).

## Conclusions

In this study we identified the emergence of a new VOI (S:E484K) within lineage B.1.1.33 circulating in Brazil. The VOI N.9 displayed a low prevalence (~3%) among all Brazilian SARS-CoV-2 samples analyzed between November 2020 and February 2021, but it is already widely dispersed in the country and comprises a high fraction (35%) of the B.1.1.33 sequences detected in that period. Mutation S:E484K has been identified as one of the most important substitutions that could contribute to immune evasion as confers resistance to several monoclonal antibodies and also reduces the neutralization potency of some polyclonal sera from convalescent and vaccinated individuals ^11-13^. Mutation S:E484K has emerged independently in multiple VOCs (P.1, B.1.351 and B.1.1.7) and VOIs (P.2 and B.1.526) ^14^ spreading around the world, and it is probably an example of convergent evolution and ongoing adaptation of the virus to the human host.

The onset date of the VOI N.9 here estimated around mid-August roughly coincides with the estimated timing of emergence of the VOI P.2 in late-July ^6^ and shortly precede the detection of a major global shift in the SARS-CoV-2 fitness landscape after October 2020 ^15^. These findings indicate that 484K variants probably arose simultaneously in the two most prevalent viral lineages circulating in Brazil around July-August, but may have only acquired some fitness advantages which accelerated its dissemination after October 2020. We predict that the Brazilian COVID-19 epidemic during 2021 will be dominated by a complex array of B.1.1.28(484K), including P.1 and P.2, and B.1.1.33(484K) variants that will completely replace the parental 484E lineages that drove the epidemic in 2020. Implementation of efficient mitigation measures in Brazil is crucial to reduce community transmission and prevent the recurrent emergence of more transmissible variants that could further exacerbate the epidemic in the country.

The opinions expressed by the authors do not necessarily reflect the opinions of the Ministry of Health of Brazil or the institutions with which the authors are affiliated.

## Acknowledgements

This study was approved by the FIOCRUZ-IOC Ethics Committee (CAAE: 68118417.6.0000.5248 and 32333120.4.0000.5190) and the Brazilian Ministry of the Environment (MMA) SISGEN (A1767C3).

The authors wish to thank all the health care workers and scientists who have worked hard to deal with this pandemic threat, the GISAID team, and all the EpiCoV database’s submitters, GISAID acknowledgment table containing sequences used in this study is attached to this post (**Appendix Table 2**). We also appreciate the support of the Fiocruz COVID-19 Genomic Surveillance Network (http://www.genomahcov.fiocruz.br/) members, the Respiratory Viruses Genomic Surveillance. General Coordination of the Laboratory Network (CGLab), Brazilian Ministry of Health (MoH), Brazilian States Central Laboratories (LACENs), Brazilian Ministry of Health (MoH), and the Amazonas surveillance teams for the partnership in the viral surveillance in Brazil. Funding support FAPEAM (PCTI-EmergeSaude/AM call 005/2020 and Rede Genômica de Vigilância em Saúde - REGESAM); Conselho Nacional de Desenvolvimento Científico e Tecnológico (grant 403276/2020-9); Inova Fiocruz/Fundação Oswaldo Cruz (Grant VPPCB-007-FIO-18-2-30 - Geração de conhecimento).

## Appendix 1.

GISAID accession numbers of genomes lineage B.1.1.33 from this study EPI_ISL_427294 to EPI_ISL_427298; EPI_ISL_427302 to EPI_ISL_427304; EPI_ISL_456071 to EPI_ISL_456077; EPI_ISL_456079 to EPI_ISL_456082; EPI_ISL_456084 to EPI_ISL_456087; EPI_ISL_456089 to EPI_ISL_456096; EPI_ISL_456098 to EPI_ISL_456106; EPI_ISL_467345; EPI_ISL_467347 to EPI_ISL_467353; EPI_ISL_467355; EPI_ISL_467357; EPI_ISL_467358; EPI_ISL_467360 to EPI_ISL_467365; EPI_ISL_467367 to EPI_ISL_467371; EPI_ISL_541347 to EPI_ISL_541350; EPI_ISL_541352; EPI_ISL_541353; EPI_ISL_541356 to EPI_ISL_541358; EPI_ISL_541360 to EPI_ISL_541365; EPI_ISL_541367; EPI_ISL_541369; EPI_ISL_541370; EPI_ISL_541376; EPI_ISL_541382; EPI_ISL_541385; EPI_ISL_541388; EPI_ISL_541392; EPI_ISL_541397 to EPI_ISL_541399; EPI_ISL_729794 to EPI_ISL_729798; EPI_ISL_729800; EPI_ISL_729802; EPI_ISL_729807; EPI_ISL_729809; EPI_ISL_729810; EPI_ISL_729812; EPI_ISL_729814; EPI_ISL_729816; EPI_ISL_729817 to EPI_ISL_729821; EPI_ISL_729823; EPI_ISL_729825; EPI_ISL_729826 to EPI_ISL_729833; EPI_ISL_729837 to EPI_ISL_729839; EPI_ISL_729841 to EPI_ISL_729844; EPI_ISL_729849; EPI_ISL_729851; EPI_ISL_729857; EPI_ISL_729858; EPI_ISL_729860; EPI_ISL_792561; EPI_ISL_792571 to EPI_ISL_792573; EPI_ISL_792579; EPI_ISL_792583; EPI_ISL_792588; EPI_ISL_792589; EPI_ISL_792593; EPI_ISL_792594; EPI_ISL_792596; EPI_ISL_792600; EPI_ISL_792601 to EPI_ISL_792603; EPI_ISL_792608; EPI_ISL_792610; EPI_ISL_792612; EPI_ISL_792623; EPI_ISL_792637; EPI_ISL_792640; EPI_ISL_792644; EPI_ISL_792648; EPI_ISL_1181353 to EPI_ISL_1181624.

**Appendix Table 1.**
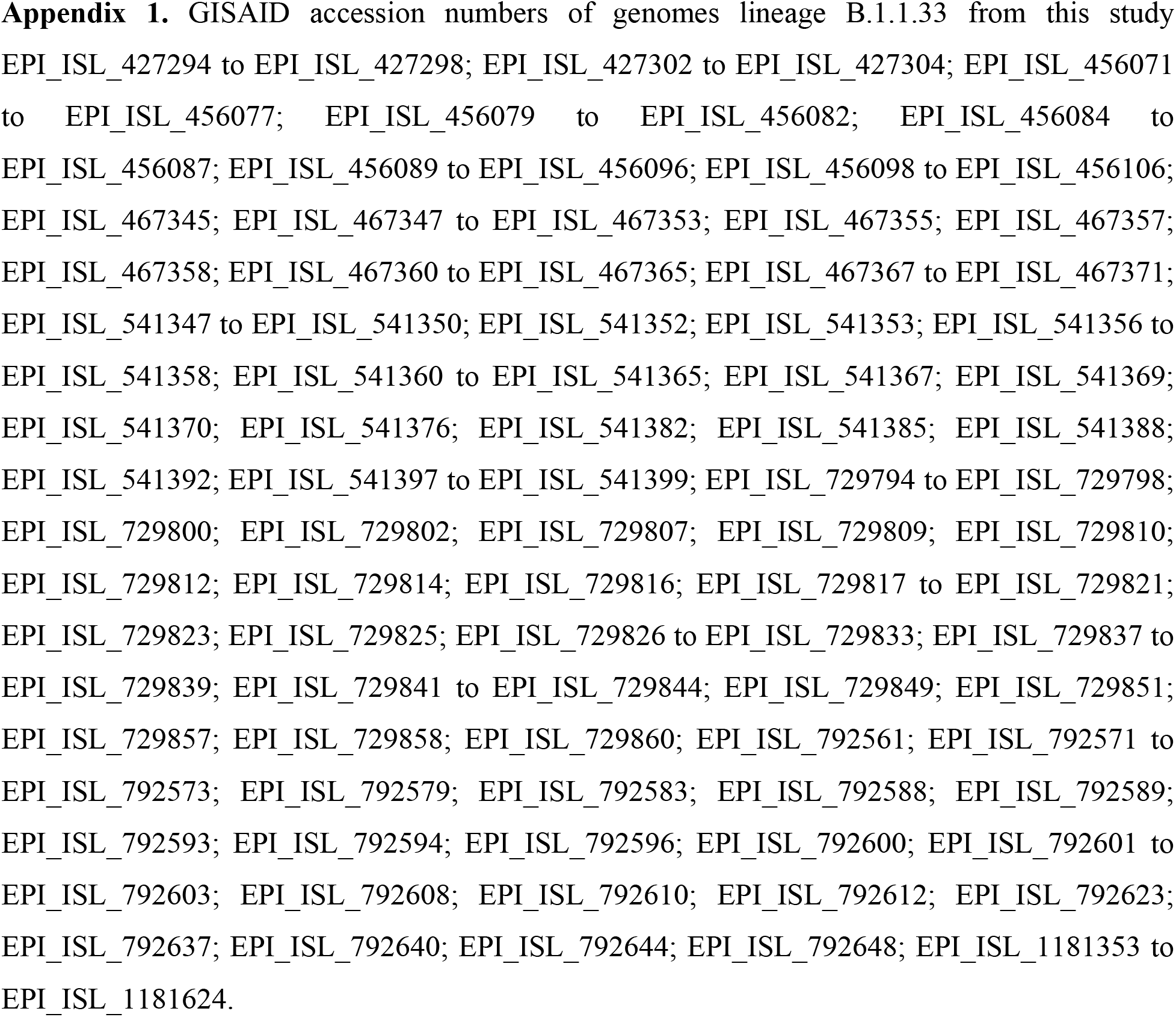
GISAID acknowledgment table.

## Notes

### Competing Interest Statement

The authors have declared no competing interest.

