## Appendix Table 1 for "A potential SARS-CoV-2 variant of interest (VOI) harboring mutation E484K in the Spike protein was identified within lineage B.1.1.33 circulating in Brazil"

We gratefully acknowledge the following Authors from the Originating laboratories responsible for obtaining the specimens, as well as the Submitting laboratories where the genome data were generated and shared via GISAID, on which this research is based.

All Submitters of data may be contacted directly via www.gisaid.org

Authors are sorted alphabetically.

| Accession ID | Originating Laboratory | Submitting Laboratory | Authors |
|---|---|---|---|
| EPI_ISL_1039700 | Instituto Adolfo Lutz Central | Instituto Adolfo Lutz, Interdiciplinary Procedures Center, Strategic Laboratory | Claudio Tavares Sacchi, Claudia Regina Gonçalves, Erica Valessa Ramos Gomes, Karoline Rodrigues Campos |
| EPI_ISL_1040824, EPI_ISL_1040829, EPI_ISL_1040831, EPI_ISL_1040833, EPI_ISL_1040835, EPI_ISL_1040836, EPI_ISL_1040837, EPI_ISL_1040839, EPI_ISL_1040840, EPI_ISL_1040842, EPI_ISL_1040843, EPI_ISL_1040844, EPI_ISL_1040845, EPI_ISL_1040848, EPI_ISL_1040851 | | | |
| see above | LACEN do Mato Grosso do Sul | Instituto Adolfo Lutz, Interdiciplinary Procedures Center, Strategic Laboratory | Claudio Tavares Sacchi, Claudia Regina Gonçalves, Erica Valessa Ramos Gomes, Karoline Rodrigues Campos |
| EPI_ISL_1060910, EPI_ISL_1060925, EPI_ISL_1060928 | DB Diagnosticos do Brasil | Instituto de Medicina Tropical de Sao Paulo | Brazil-UK Centre for Arbovirus Discovery Diagnosis Genomics and Epidemiology (CADDE) Genomic Network - Instituto de Medicina Tropical |
| EPI_ISL_1060934, EPI_ISL_1060983, EPI_ISL_1060988 | CDL Laboratorio Santos e Vidal LTDA. | Instituto de Medicina Tropical de Sao Paulo | Brazil-UK Centre for Arbovirus Discovery Diagnosis Genomics and Epidemiology (CADDE) Genomic Network - Instituto de Medicina Tropical |
| EPI_ISL_1060994 | DB Diagnosticos do Brasil | Instituto de Medicina Tropical de Sao Paulo | Brazil-UK Centre for Arbovirus Discovery Diagnosis Genomics and Epidemiology (CADDE) Genomic Network - Instituto de Medicina Tropical |
| EPI_ISL_1061020, EPI_ISL_1064743 | CDL Laboratorio Santos e Vidal LTDA. | Instituto de Medicina Tropical de Sao Paulo | Brazil-UK Centre for Arbovirus Discovery Diagnosis Genomics and Epidemiology (CADDE) Genomic Network - Instituto de Medicina Tropical |
| EPI_ISL_1068083, EPI_ISL_1068094, EPI_ISL_1068097, EPI_ISL_1068098, EPI_ISL_1068099, EPI_ISL_1068103, EPI_ISL_1068120, EPI_ISL_1068121, EPI_ISL_1068122, EPI_ISL_1068139, EPI_ISL_1068140, EPI_ISL_1068144, EPI_ISL_1068163, EPI_ISL_1068189, EPI_ISL_1068200, EPI_ISL_1068202, EPI_ISL_1068204, EPI_ISL_1068216, EPI_ISL_1068220, EPI_ISL_1068228, EPI_ISL_1068229, EPI_ISL_1068230, EPI_ISL_1068234, EPI_ISL_1068240, EPI_ISL_1068241, EPI_ISL_1068255, EPI_ISL_1068265 | | | |
| see above | Laboratorio de Ecologia de Doencas Transmissiveis na Amazonia, Instituto Leonidas e Maria Deane - Fiocruz Amazonia | Laboratorio de Ecologia de Doencas Transmissiveis na Amazonia, Instituto Leonidas e Maria Deane - Fiocruz Amazonia | Valdinete Nascimento, Victor Souza, André Corado, Fernanda Nascimento, George Silva, Ágatha Costa, Debora Duarte, Karina Pessoa, Matilde Mejía, Luciana Gonçalves, Maria Júlia Brandão, Michele Jesus, Felipe Naveca |
| EPI_ISL_1068315, EPI_ISL_1068316, EPI_ISL_1068317, EPI_ISL_1068318, EPI_ISL_1068320, EPI_ISL_1068321, EPI_ISL_1068323, EPI_ISL_1068324, EPI_ISL_1068326, EPI_ISL_1068328, EPI_ISL_1068331, EPI_ISL_1068332, EPI_ISL_1068334, EPI_ISL_1068336, EPI_ISL_1068337, EPI_ISL_1068339, EPI_ISL_1068342, EPI_ISL_1068343, EPI_ISL_1068345, EPI_ISL_1068366, EPI_ISL_1068367, EPI_ISL_1068372, EPI_ISL_1068374, EPI_ISL_1068375, EPI_ISL_1068379, EPI_ISL_1068382, EPI_ISL_1068383, EPI_ISL_1068385, EPI_ISL_1068386, EPI_ISL_1068387, EPI_ISL_1068388, EPI_ISL_1068390, EPI_ISL_1068391, EPI_ISL_1068392, EPI_ISL_1068393 | | | |
| see above | Central Public Health Laboratory - LACEN -Bahia, Salvador, Brazil | Central Public Health Laboratory - LACEN -Bahia, Salvador, Brazil | Stephane Tosta, Luciana Oliveira, Vanessa Nardy,Patrícia Cajado,Marcela Gómez, Breno Dominguez, Jaqueline Gomes, Vagner Fonseca,Marta Giovanetti,Luiz Alcantara, Felicidade Pereira, Arabela Leal |
| EPI_ISL_1079159 | IAL Regional de Bauru | Instituto Adolfo Lutz, Interdiciplinary Procedures Center, Strategic Laboratory | Claudio Tavares Sacchi, Claudia Regina Gonçalves, Erica Valessa Ramos Gomes, Karoline Rodrigues Campos |
| EPI_ISL_1084726 | DB Diagnosticos do Brasil | Laboratório de Parasitologia Médica - Instituto de Medicina Tropical - Universidade de São Paulo | Brazil-UK Centre for Arbovirus Discovery Diagnosis Genomics and Epidemiology (CADDE) Genomic Network - Instituto de Medicina Tropical |
| EPI_ISL_1084729 | IMT_USP | Laboratório de Parasitologia Médica - Instituto de Medicina Tropical - Universidade de São Paulo | Brazil-UK Centre for Arbovirus Discovery Diagnosis Genomics and Epidemiology (CADDE) Genomic Network - Instituto de Medicina Tropical |
| EPI_ISL_1084730, EPI_ISL_1084740 | DB Diagnosticos do Brasil | Laboratório de Parasitologia Médica - Instituto de Medicina Tropical - Universidade de São Paulo | Brazil-UK Centre for Arbovirus Discovery Diagnosis Genomics and Epidemiology (CADDE) Genomic Network - Instituto de Medicina Tropical |
| EPI_ISL_402124 | Wuhan Jinyintan Hospital | Wuhan Institute of Virology, Chinese Academy of Sciences | Peng Zhou, Xing-Lou Yang, Ding-Yu Zhang, Lei Zhang, Yan Zhu, Hao-Rui Si, Zhengli Shi |
| EPI_ISL_427294, EPI_ISL_427295, EPI_ISL_427296, EPI_ISL_427297, EPI_ISL_427298, EPI_ISL_427302, EPI_ISL_427303, EPI_ISL_427304 | Instituto Oswaldo Cruz FIOCRUZ - Laboratory of Respiratory Viruses and Measles (LVRS) | Instituto Oswaldo Cruz FIOCRUZ - Laboratory of Respiratory Viruses and Measles (LVRS) | Paola Resende, Fernando Motta, Luciana Appolinario, Sunando Roy, Aline Mattos, Milene Miranda, Cristiana Garcia, Braulia Caetano, Maria Ogrzewalska, Priscila Born, Jonathan Lopes, Marilda Siqueira |
| EPI_ISL_450873 | Evandro Chagas Institute | Evandro Chagas Institute | Santos, M.C.; Silva, A.M.; Junior, W.D.C.; Barbagelata, L.S.; Ferreira, J.A.; Sousa, E.M.A.; da Silva, P.S.; Martins, L.C.; Sousa Junior, E.C.; Viana, G.M.R |
| EPI_ISL_450874 | Evandro Chagas Institute | Evandro Chagas Institute | Santos, M.C.; Silva, A.M.; Junior, W.D.C.; Barbagelata, L.S.; Ferreira, J.A.; Sousa, E.M.A.; da Silva, P.S.; Martins, L.C.;Sousa Junior, E.C.;Viana, G.M.R |
| EPI_ISL_456071, EPI_ISL_456072, EPI_ISL_456073, EPI_ISL_456074, EPI_ISL_456075 | Laboratory of Respiratory Viruses and Measles, Oswaldo Cruz Institute, FIOCRUZ | Laboratory of Respiratory Viruses and Measles, Oswaldo Cruz Institute, FIOCRUZ | Paola Resende, Luciana Appolinario, Fernando Motta, Aline Mattos, Milene Miranda, Cristiana Garcia, Braulia Caetano, Maria Ogrzewalska, Jonathan Lopes, Marilda Siqueira |
| EPI_ISL_456076, EPI_ISL_456077 | LACEN RJ - Laboratório Central de Saúde Pública Noel Nutels | Laboratory of Respiratory Viruses and Measles, Oswaldo Cruz Institute, FIOCRUZ | Paola Resende, Luciana Appolinario, Fernando Motta, Aline Mattos, Milene Miranda, Cristiana Garcia, Braulia Caetano, Maria Ogrzewalska, Jonathan Lopes, Marilda Siqueira |
| EPI_ISL_456079, EPI_ISL_456080, EPI_ISL_456081 | Laboratory of Respiratory Viruses and Measles, Oswaldo Cruz Institute, FIOCRUZ | Laboratory of Respiratory Viruses and Measles, Oswaldo Cruz Institute, FIOCRUZ | Paola Resende, Luciana Appolinario, Fernando Motta, Aline Mattos, Milene Miranda, Cristiana Garcia, Braulia Caetano, Maria Ogrzewalska, Jonathan Lopes, Marilda Siqueira |
| EPI_ISL_456082 | LACEN RJ - Laboratório Central de Saúde Pública Noel Nutels | Laboratory of Respiratory Viruses and Measles, Oswaldo Cruz Institute, FIOCRUZ | Paola Resende, Luciana Appolinario, Fernando Motta, Aline Mattos, Milene Miranda, Cristiana Garcia, Braulia Caetano, Maria Ogrzewalska, Jonathan Lopes, Marilda Siqueira |
| EPI_ISL_456084, EPI_ISL_456085, EPI_ISL_456086, EPI_ISL_456087, EPI_ISL_456089, EPI_ISL_456090, EPI_ISL_456091, EPI_ISL_456092, EPI_ISL_456093, EPI_ISL_456094, EPI_ISL_456095, EPI_ISL_456096, EPI_ISL_456098, EPI_ISL_456099, EPI_ISL_456100, EPI_ISL_456101, EPI_ISL_456102, EPI_ISL_456103, EPI_ISL_456104, EPI_ISL_456105, EPI_ISL_456106 | | | |
| see above | Laboratory of Respiratory Viruses and Measles, Oswaldo Cruz Institute, FIOCRUZ | Laboratory of Respiratory Viruses and Measles, Oswaldo Cruz Institute, FIOCRUZ | Paola Resende, Luciana Appolinario, Fernando Motta, Aline Mattos, Milene Miranda, Cristiana Garcia, Braulia Caetano, Maria Ogrzewalska, Jonathan Lopes, Marilda Siqueira |
| EPI_ISL_458138, EPI_ISL_458139, EPI_ISL_458142, EPI_ISL_458143, EPI_ISL_458144, EPI_ISL_458145, EPI_ISL_458148, EPI_ISL_458149 | Evandro Chagas Institute | Evandro Chagas Institute | Santos, M.C.; Silva, A.M.; Junior, W.D.C.; Barbagelata, L.S.; Ferreira, J.A.; Sousa, E.M.A.; da Silva, P.S.; Resque, H.R; Martins, L.C.; Sousa Junior, E.C.;Viana, G.M.R |
| EPI_ISL_467345, EPI_ISL_467347, EPI_ISL_467348, EPI_ISL_467349, EPI_ISL_467350, EPI_ISL_467351, EPI_ISL_467352, EPI_ISL_467353, EPI_ISL_467355, EPI_ISL_467357, EPI_ISL_467358, EPI_ISL_467360, EPI_ISL_467361, EPI_ISL_467362, EPI_ISL_467363, EPI_ISL_467364, EPI_ISL_467365, EPI_ISL_467367, EPI_ISL_467368, EPI_ISL_467369, EPI_ISL_467370, EPI_ISL_467371 | | | |
| see above | Laboratory of Respiratory Viruses and Measles, Oswaldo Cruz Institute, FIOCRUZ | Laboratory of Respiratory Viruses and Measles, Oswaldo Cruz Institute, FIOCRUZ | Paola Resende, Luciana Appolinario, Fernando Motta, Anna Carolina Paixão, Ana Carolina Mendonça, Aline Mattos, Milene Miranda, Cristiana Garcia, Braulia Caetano, Maria Ogrzewalska, Jonathan Lopes, Marilda Siqueira |
| EPI_ISL_468310 | Hospital Sao Paulo de Ensino da UNIFESP | Instituto Adolfo Lutz, Interdiciplinary Procedures Center, Strategic Laboratory | Claudio Tavares Sacchi, Claudia Regina Gonçalves, Erica Valessa Ramos Gomes |
| EPI_ISL_470570, EPI_ISL_470572, EPI_ISL_470577, EPI_ISL_470580, EPI_ISL_470582, EPI_ISL_470583, EPI_ISL_470584, EPI_ISL_470585, EPI_ISL_470586, EPI_ISL_470587, EPI_ISL_470588 | | | |

| | | | |
|---|---|---|---|
| see above | Hermes Pardini | Bioinformatics Laboratory / LNCC | Alexandra Gerber, Ana Paula Guimarães, Luiz Gonzaga Paula de Almeida, Ronaldo da Silva Francisco Junior, Mariane Talon, Filipe Romero, Átila Duque Rossi, Terezinha Marta Pereira, working group UFRJ, Jaqueline Goes de Jesus, Ingra Morales Claro, Ester Cerdeira Sabino, Nuno Rodrigues Faria, CADDE-group, Laboratorio Hermes Pardini, Laboratorio Simile, working group UFMG, Amilcar Tanuri, Carolina Voloch, Renato Santana Aguiar e Ana Tereza Vasconcelos |
| EPI_ISL_470589, EPI_ISL_470591, EPI_ISL_470592, EPI_ISL_470594, EPI_ISL_470595, EPI_ISL_470596, EPI_ISL_470597 | Simile | Bioinformatics Laboratory / LNCC | Alexandra Gerber, Ana Paula Guimarães, Luiz Gonzaga Paula de Almeida, Ronaldo da Silva Francisco Junior, Mariane Talon, Filipe Romero, Átila Duque Rossi, Terezinha Marta Pereira, working group UFRJ, Jaqueline Goes de Jesus, Ingra Morales Claro, Ester Cerdeira Sabino, Nuno Rodrigues Faria, CADDE-group, Laboratorio Hermes Pardini, Laboratorio Simile, working group UFMG, Amilcar Tanuri, Carolina Voloch, Renato Santana Aguiar e Ana Tereza Vasconcelos |
| EPI_ISL_470601, EPI_ISL_470606, EPI_ISL_470609, EPI_ISL_470611, EPI_ISL_470613, EPI_ISL_470614 | Hermes Pardini | Bioinformatics Laboratory / LNCC | Alexandra Gerber, Ana Paula Guimarães, Luiz Gonzaga Paula de Almeida, Ronaldo da Silva Francisco Junior, Mariane Talon, Filipe Romero, Átila Duque Rossi, Terezinha Marta Pereira, working group UFRJ, Jaqueline Goes de Jesus, Ingra Morales Claro, Ester Cerdeira Sabino, Nuno Rodrigues Faria, CADDE-group, Laboratorio Hermes Pardini, Laboratorio Simile, working group UFMG, Amilcar Tanuri, Carolina Voloch, Renato Santana Aguiar e Ana Tereza Vasconcelos |
| EPI_ISL_470615, EPI_ISL_470617, EPI_ISL_470618, EPI_ISL_470620, EPI_ISL_470621, EPI_ISL_470623, EPI_ISL_470624, EPI_ISL_470625, EPI_ISL_470626, EPI_ISL_470627, EPI_ISL_470628, EPI_ISL_470629, EPI_ISL_470630, EPI_ISL_470631, EPI_ISL_470632, EPI_ISL_470633, EPI_ISL_470634, EPI_ISL_470635, EPI_ISL_470636, EPI_ISL_470637, EPI_ISL_470639, EPI_ISL_470640, EPI_ISL_470641, EPI_ISL_470642, EPI_ISL_470643, EPI_ISL_470644, EPI_ISL_470645, EPI_ISL_470646, EPI_ISL_470647, EPI_ISL_470648, EPI_ISL_470649, EPI_ISL_470650 | | | |
| see above | Laboratorio de Virologia Molecular / UFRJ | Bioinformatics Laboratory / LNCC | Alexandra Gerber, Ana Paula Guimarães, Luiz Gonzaga Paula de Almeida, Ronaldo da Silva Francisco Junior, Mariane Talon, Filipe Romero, Átila Duque Rossi, Terezinha Marta Pereira, working group UFRJ, Jaqueline Goes de Jesus, Ingra Morales Claro, Ester Cerdeira Sabino, Nuno Rodrigues Faria, CADDE-group, Laboratorio Hermes Pardini, Laboratorio Simile, working group UFMG, Amilcar Tanuri, Carolina Voloch, Renato Santana Aguiar e Ana Tereza Vasconcelos |
| EPI_ISL_470655 | Hermes Pardini | Bioinformatics Laboratory / LNCC | Alexandra Gerber, Ana Paula Guimarães, Luiz Gonzaga Paula de Almeida, Ronaldo da Silva Francisco Junior, Mariane Talon, Filipe Romero, Átila Duque Rossi, Terezinha Marta Pereira, working group UFRJ, Jaqueline Goes de Jesus, Ingra Morales Claro, Ester Cerdeira Sabino, Nuno Rodrigues Faria, CADDE-group, Laboratorio Hermes Pardini, Laboratorio Simile, working group UFMG, Amilcar Tanuri, Carolina Voloch, Renato Santana Aguiar e Ana Tereza Vasconcelos |
| EPI_ISL_471551 | Hospital Sao Paulo de Ensino da Unifesp | Instituto Adolfo Lutz, Interdiciplinary Procedures Center, Strategic Laboratory | Claudio Tavares Sacchi, Claudia Regina Gonçalves, Erica Valessa Ramos Gomes |
| EPI_ISL_471554 | Hospital Bosque da Saúde | Instituto Adolfo Lutz, Interdiciplinary Procedures Center, Strategic Laboratory | Claudio Tavares Sacchi, Claudia Regina Gonçalves, Erica Valessa Ramos Gomes |
| EPI_ISL_476155, EPI_ISL_476158, EPI_ISL_476160, EPI_ISL_476168, EPI_ISL_476170 | Laboratório de Patologia Clínica - UNICAMP | Laboratório de Estudos de Vírus Emergentes - UNICAMP | José Luiz Proença-Modena, Magnun Nueldo Nunes dos Santos, Angelica Schreiber, Julia Forato,Camila Simeoni, Marcilio Jorge Fumagalli, Mariene Ribeiro Amorim, Darlan da Silva Candido, Nuno Rodrigues Faria, Julien Theze, Luiz Gonzaga,Jaqueline Goes Jesus e William Marciel de Souza |
| EPI_ISL_476171, EPI_ISL_476172, EPI_ISL_476173, EPI_ISL_476174, EPI_ISL_476178, EPI_ISL_476180, EPI_ISL_476188, EPI_ISL_476189, EPI_ISL_476190, EPI_ISL_476191, EPI_ISL_476192, EPI_ISL_476194, EPI_ISL_476195, EPI_ISL_476201, EPI_ISL_476202 | | | |
| see above | DB Diagnósticos do Brasil | Instituto de Medicina Tropical da Univesidade de São Paulo | Samples: Nelson Gaburo Jr; Sequencing: Ingra Morales Claro, Jaqueline Goes de Jesus, Erika Regina Manuli, Flavia Cristina da Silva Sales, Thais de Moura Coletti, Camila Alves Maia da Silva, Mariana Severo Ramundo, Giula Magalhaes Ferreira, Darlan da Silva Candido, Julien Theze, Nuno Faria, Ester Sabino |
| EPI_ISL_476204 | Hospital da Clínicas da Faculdade de Medicina da Universidade de São Paulo | Instituto de Medicina Tropical da Univesidade de São Paulo | Samples: Ingra Morales Claro, Erika Regina Manuli, Cecilia Salete Alencar, Carolina S. Lazar, Silvia F. Costa; Sequencing: Ingra Morales Claro, Jaqueline Goes de Jesus, Erika Regina Manuli, Flavia Cristina da Silva Sales, Thais de Moura Coletti, Camila Alves Maia da Silva, Mariana Severo Ramundo, Giula Magalhaes Ferreira, Darlan da Silva Candido, Julien Theze, Nuno Faria, Ester Sabino |
| EPI_ISL_476210, EPI_ISL_476211, EPI_ISL_476215, EPI_ISL_476216 | DB Diagnósticos do Brasil | Instituto de Medicina Tropical da Univesidade de São Paulo | Samples: Nelson Gaburo Jr; Sequencing: Ingra Morales Claro, Jaqueline Goes de Jesus, Erika Regina Manuli, Flavia Cristina da Silva Sales, Thais de Moura Coletti, Camila Alves Maia da Silva, Mariana Severo Ramundo, Giula Magalhaes Ferreira, Darlan da Silva Candido, Julien Theze, Nuno Faria, Ester Sabino |
| EPI_ISL_476221 | Laboratory Fleury | Instituto de Medicina Tropical da Univesidade de São Paulo | Samples: Celso Granato; Sequencing: Ingra Morales Claro, Jaqueline Goes de Jesus, Erika Regina Manuli, Flavia Cristina da Silva Sales, Thais de Moura Coletti, Camila Alves Maia da Silva, Mariana Severo Ramundo, Giula Magalhaes Ferreira, Darlan da Silva Candido, Julien Theze, Nuno Faria, Ester Sabino |
| EPI_ISL_476245, EPI_ISL_476246, EPI_ISL_476247, EPI_ISL_476248, EPI_ISL_476263, EPI_ISL_476273, EPI_ISL_476274, EPI_ISL_476276 | Hospital da Clínicas da Faculdade de Medicina da Universidade de São Paulo | Instituto de Medicina Tropical da Univesidade de São Paulo | Samples: Ingra Morales Claro, Erika Regina Manuli, Cecilia Salete Alencar, Carolina S. Lazar, Silvia F. Costa; Sequencing: Ingra Morales Claro, Jaqueline Goes de Jesus, Erika Regina Manuli, Flavia Cristina da Silva Sales, Thais de Moura Coletti, Camila Alves Maia da Silva, Mariana Severo Ramundo, Giula Magalhaes Ferreira, Darlan da Silva Candido, Julien Theze, Nuno Faria, Ester Sabino |
| EPI_ISL_476278, EPI_ISL_476281, EPI_ISL_476286, EPI_ISL_476290, EPI_ISL_476292, EPI_ISL_476295, EPI_ISL_476297, EPI_ISL_476300, EPI_ISL_476301, EPI_ISL_476302, EPI_ISL_476303, EPI_ISL_476305, EPI_ISL_476309, EPI_ISL_476310, EPI_ISL_476315, EPI_ISL_476316, EPI_ISL_476317, EPI_ISL_476319, EPI_ISL_476324, EPI_ISL_476327, EPI_ISL_476328, EPI_ISL_476329, EPI_ISL_476330, EPI_ISL_476332, EPI_ISL_476334, EPI_ISL_476335, EPI_ISL_476350, EPI_ISL_476351, EPI_ISL_476352, EPI_ISL_476355, EPI_ISL_476356, EPI_ISL_476358, EPI_ISL_476360, EPI_ISL_476362, EPI_ISL_476365 | | | |
| see above | DB Diagnósticos do Brasil | Instituto de Medicina Tropical da Univesidade de São Paulo | Samples: Nelson Gaburo Jr; Sequencing: Ingra Morales Claro, Jaqueline Goes de Jesus, Erika Regina Manuli, Flavia Cristina da Silva Sales, Thais de Moura Coletti, Camila Alves Maia da Silva, Mariana Severo Ramundo, Giula Magalhaes Ferreira, Darlan da Silva Candido, Julien Theze, Nuno Faria, Ester Sabino |
| EPI_ISL_476374, EPI_ISL_476377, EPI_ISL_476383 | Hospital da Clínicas da Faculdade de Medicina da Universidade de São Paulo | Instituto de Medicina Tropical da Univesidade de São Paulo | Samples: Ingra Morales Claro, Erika Regina Manuli, Cecilia Salete Alencar, Carolina S. Lazar, Silvia F. Costa; Sequencing: Ingra Morales Claro, Jaqueline Goes de Jesus, Erika Regina Manuli, Flavia Cristina da Silva Sales, Thais de Moura Coletti, Camila Alves Maia da Silva, Mariana Severo Ramundo, Giula Magalhaes Ferreira, Darlan da Silva Candido, Julien Theze, Nuno Faria, Ester Sabino |
| EPI_ISL_476392, EPI_ISL_476422, EPI_ISL_476425 | Laboratório de Patologia Clínica - UNICAMP | Laboratório de Estudos de Vírus Emergentes - UNICAMP | José Luiz Proença-Modena, Magnun Nueldo Nunes dos Santos, Angelica Schreiber, Julia Forato,Camila Simeoni, Marcilio Jorge Fumagalli, Mariene Ribeiro Amorim, Darlan da Silva Candido, Nuno Rodrigues Faria, Julien Theze, Luiz Gonzaga,Jaqueline Goes Jesus e William Marciel de Souza |
| EPI_ISL_476429, EPI_ISL_476435, EPI_ISL_476439, EPI_ISL_476444, EPI_ISL_476449, EPI_ISL_476450, EPI_ISL_476459, EPI_ISL_476483, EPI_ISL_476484, EPI_ISL_476486, EPI_ISL_476487 | | | |
| see above | Hospital da Clínicas da Faculdade de Medicina da Universidade de São Paulo | Instituto de Medicina Tropical da Univesidade de São Paulo | Samples: Ingra Morales Claro, Erika Regina Manuli, Cecilia Salete Alencar, Carolina S. Lazar, Silvia F. Costa; Sequencing: Ingra Morales Claro, Jaqueline Goes de Jesus, Erika Regina Manuli, Flavia Cristina da Silva Sales, Thais de Moura Coletti, Camila Alves Maia da Silva, Mariana Severo Ramundo, Giula Magalhaes Ferreira, Darlan da Silva Candido, Julien Theze, Nuno Faria, Ester Sabino |
| EPI_ISL_486427 | unknown | Clinical Laboratory, Hospital Israelita Albert Einstein | Amgarteb,D., Malta,F., Guedes,R.L., Santana,R.A., de Menezes,F.G., Mangueira,C.L. and Pinho,J.R. |
| EPI_ISL_492032 | Instituto de Biologia do Exército | Laboratório Metabolismo Macromolecular FirminoTorres de Castro, Instituto de Biofísica Carlos Chagas Filho, Universidade Federal do Rio de Janeiro | Bianca Catarina Azevedo Cabral, Aline Rosa Vianna de Souza , Marcos Dornelas-Ribeiro, Tatiana LS Nogueira, Nádia Vaez Gonçalves da Cruz, Caleb GM Santos, Elizabeth Valentin, Marcio da Costa Cipitelli, Virginia Sara Grancieri do Amaral, Rodrigo Soares de Moura Neto, Clarissa Damaso, Rosane Silva |
| EPI_ISL_492033 | Instituto de Biologia do Exército | Laboratório Metabolismo Macromolecular FirminoTorres de Castro, Instituto de Biofísica Carlos Chagas Filho, Universidade Federal do Rio de Janeiro | Bianca Catarina Azevedo Cabral, Aline Rosa Vianna de Souza, Caleb GM Santos, Marcos Dornelas-Ribeiro, Tatiana LS Nogueira, Nádia Vaez Gonçalves da Cruz, Elizabeth Valentin, Marcio da Costa Cipitelli, Virginia Sara Grancieri do Amaral, Rodrigo Soares de Moura Neto, Clarissa Damaso, Rosane Silva |
| EPI_ISL_492034 | Instituto de Biologia do Exército | Laboratório Metabolismo Macromolecular FirminoTorres de Castro, Instituto de Biofísica Carlos Chagas Filho, Universidade Federal do Rio de Janeiro | Bianca Catarina Azevedo Cabral, Aline Rosa Vianna de Souza, Nádia Vaez Gonçalves da Cruz, Caleb GM Santos, Marcos Dornelas-Ribeiro, Tatiana LS Nogueira, Elizabeth Valentin, Marcio da Costa Cipitelli, Virginia Sara Grancieri do Amaral, Rodrigo Soares de Moura Neto, Clarissa Damaso, Rosane Silva |
| EPI_ISL_492035 | Instituto de Biologia do Exército | Laboratório Metabolismo Macromolecular FirminoTorres de Castro, Instituto de Biofísica Carlos Chagas Filho, Universidade Federal do Rio de Janeiro | Bianca Catarina Azevedo Cabral, Aline Rosa Vianna de Souza, Tatiana LS Nogueira, Nádia Vaez Gonçalves da Cruz, Caleb GM Santos, Marcos Dornelas-Ribeiro, Elizabeth Valentin, Marcio da Costa Cipitelli, Virginia Sara Grancieri do Amaral, Rodrigo Soares de Moura Neto, Clarissa Damaso, Rosane Silva |
| EPI_ISL_492037 | Instituto de Biologia do Exército | Laboratório Metabolismo Macromolecular FirminoTorres de | Bianca Catarina Azevedo Cabral, Aline Rosa Vianna de Souza, Caleb GM Santos, Marcos Dornelas-Ribeiro, Tatiana LS Nogueira, Nádia Vaez Gonçalves |

| | | | |
|---|---|---|---|
| | | Castro, Instituto de Biofísica Carlos Chagas Filho, Universidade Federal do Rio de Janeiro | da Cruz, Elizabeth Valentin, Marcio da Costa Cipitelli, Virginia Sara Grancieri do Amaral, Rodrigo Soares de Moura Neto, Clarissa Damaso, Rosane Silva |
| EPI_ISL_492038 | Instituto de Biologia do Exército | Laboratório Metabolismo Macromolecular FirminoTorres de Castro, Instituto de Biofísica Carlos Chagas Filho, Universidade Federal do Rio de Janeiro | Bianca Catarina Azevedo Cabral, Aline Rosa Vianna de Souza, Nádia Vaez Gonçalves da Cruz, Caleb GM Santos, Marcos Dornelas-Ribeiro, Tatiana LS Nogueira, Elizabeth Valentin, Marcio da Costa Cipitelli, Virginia Sara Grancieri do Amaral, Rodrigo Soares de Moura Neto, Clarissa Damaso, Rosane Silva |
| EPI_ISL_492039 | Instituto de Biologia do Exército | Laboratório Metabolismo Macromolecular FirminoTorres de Castro, Instituto de Biofísica Carlos Chagas Filho, Universidade Federal do Rio de Janeiro | Bianca Catarina Azevedo Cabral, Aline Rosa Vianna de Souza, Tatiana LS Nogueira, Nádia Vaez Gonçalves da Cruz, Caleb GM Santos, Marcos Dornelas-Ribeiro, Elizabeth Valentin, Marcio da Costa Cipitelli, Virginia Sara Grancieri do Amaral, Rodrigo Soares de Moura Neto, Clarissa Damaso, Rosane Silva |
| EPI_ISL_492040 | Instituto de Biologia do Exército | Laboratório Metabolismo Macromolecular FirminoTorres de Castro, Instituto de Biofísica Carlos Chagas Filho, Universidade Federal do Rio de Janeiro | Bianca Catarina Azevedo Cabral, Aline Rosa Vianna de Souza , Marcos Dornelas-Ribeiro, Tatiana LS Nogueira, Nádia Vaez Gonçalves da Cruz, Caleb GM Santos, Elizabeth Valentin, Marcio da Costa Cipitelli, Virginia Sara Grancieri do Amaral, Rodrigo Soares de Moura Neto, Clarissa Damaso, Rosane Silva |
| EPI_ISL_492041 | Instituto de Biologia do Exército | Laboratório Metabolismo Macromolecular FirminoTorres de Castro, Instituto de Biofísica Carlos Chagas Filho, Universidade Federal do Rio de Janeiro | Bianca Catarina Azevedo Cabral, Aline Rosa Vianna de Souza, Caleb GM Santos, Marcos Dornelas-Ribeiro, Tatiana LS Nogueira, Nádia Vaez Gonçalves da Cruz, Elizabeth Valentin, Marcio da Costa Cipitelli, Virginia Sara Grancieri do Amaral, Rodrigo Soares de Moura Neto, Clarissa Damaso, Rosane Silva |
| EPI_ISL_492042 | Instituto de Biologia do Exército | Laboratório Metabolismo Macromolecular FirminoTorres de Castro, Instituto de Biofísica Carlos Chagas Filho, Universidade Federal do Rio de Janeiro | Bianca Catarina Azevedo Cabral, Aline Rosa Vianna de Souza, Nádia Vaez Gonçalves da Cruz, Caleb GM Santos, Marcos Dornelas-Ribeiro, Tatiana LS Nogueira, Elizabeth Valentin, Marcio da Costa Cipitelli, Virginia Sara Grancieri do Amaral, Rodrigo Soares de Moura Neto, Clarissa Damaso, Rosane Silva |
| EPI_ISL_492043 | Instituto de Biologia do Exército | Laboratório Metabolismo Macromolecular FirminoTorres de Castro, Instituto de Biofísica Carlos Chagas Filho, Universidade Federal do Rio de Janeiro | Bianca Catarina Azevedo Cabral, Aline Rosa Vianna de Souza, Tatiana LS Nogueira, Nádia Vaez Gonçalves da Cruz, Caleb GM Santos, Marcos Dornelas-Ribeiro, Elizabeth Valentin, Marcio da Costa Cipitelli, Virginia Sara Grancieri do Amaral, Rodrigo Soares de Moura Neto, Clarissa Damaso, Rosane Silva |
| EPI_ISL_492044 | Instituto de Biologia do Exército | Laboratório Metabolismo Macromolecular FirminoTorres de Castro, Instituto de Biofísica Carlos Chagas Filho, Universidade Federal do Rio de Janeiro | Bianca Catarina Azevedo Cabral, Aline Rosa Vianna de Souza , Marcos Dornelas-Ribeiro, Tatiana LS Nogueira, Nádia Vaez Gonçalves da Cruz, Caleb GM Santos, Elizabeth Valentin, Marcio da Costa Cipitelli, Virginia Sara Grancieri do Amaral, Rodrigo Soares de Moura Neto, Clarissa Damaso, Rosane Silva |
| EPI_ISL_492045 | Instituto de Biologia do Exército | Laboratório Metabolismo Macromolecular FirminoTorres de Castro, Instituto de Biofísica Carlos Chagas Filho, Universidade Federal do Rio de Janeiro | Bianca Catarina Azevedo Cabral, Aline Rosa Vianna de Souza, Caleb GM Santos, Marcos Dornelas-Ribeiro, Tatiana LS Nogueira, Nádia Vaez Gonçalves da Cruz, Elizabeth Valentin, Marcio da Costa Cipitelli, Virginia Sara Grancieri do Amaral, Rodrigo Soares de Moura Neto, Clarissa Damaso, Rosane Silva |
| EPI_ISL_492046 | Instituto de Biologia do Exército | Laboratório Metabolismo Macromolecular FirminoTorres de Castro, Instituto de Biofísica Carlos Chagas Filho, Universidade Federal do Rio de Janeiro | Bianca Catarina Azevedo Cabral, Aline Rosa Vianna de Souza, Nádia Vaez Gonçalves da Cruz, Caleb GM Santos, Marcos Dornelas-Ribeiro, Tatiana LS Nogueira, Elizabeth Valentin, Marcio da Costa Cipitelli, Virginia Sara Grancieri do Amaral, Rodrigo Soares de Moura Neto, Clarissa Damaso, Rosane Silva |
| EPI_ISL_492047 | Instituto de Biologia do Exército | Laboratório Metabolismo Macromolecular FirminoTorres de Castro, Instituto de Biofísica Carlos Chagas Filho, Universidade Federal do Rio de Janeiro | Bianca Catarina Azevedo Cabral, Aline Rosa Vianna de Souza, Tatiana LS Nogueira, Nádia Vaez Gonçalves da Cruz, Caleb GM Santos, Marcos Dornelas-Ribeiro, Elizabeth Valentin, Marcio da Costa Cipitelli, Virginia Sara Grancieri do Amaral, Rodrigo Soares de Moura Neto, Clarissa Damaso, Rosane Silva |
| EPI_ISL_492048 | Instituto de Biologia do Exército | Laboratório Metabolismo Macromolecular FirminoTorres de Castro, Instituto de Biofísica Carlos Chagas Filho, Universidade Federal do Rio de Janeiro | Bianca Catarina Azevedo Cabral, Aline Rosa Vianna de Souza , Marcos Dornelas-Ribeiro, Tatiana LS Nogueira, Nádia Vaez Gonçalves da Cruz, Caleb GM Santos, Elizabeth Valentin, Marcio da Costa Cipitelli, Virginia Sara Grancieri do Amaral, Rodrigo Soares de Moura Neto, Clarissa Damaso, Rosane Silva |
| EPI_ISL_500479 | LACEN/PE | WallauLab, Aggeu Magalhaes Institute | Marcelo Henrique Santos Paiva, Duschinka Ribeiro Duarte Guedes, Cássia Docena, Matheus Filgueira Bezerra, Filipe Zimmer Dezordi, Laís Ceschini Machado, Larissa Krokovsky, Elisama Helvecio, Alexandre Freitas da Silva, Luydson Richardson Silva Vasconcelos, Antonio Mauro Rezende, Severino Jefferson Ribeiro da Silva, Kamila Gaudêncio da Silva Sales, Bruna Santos Lima Figueiredo de Sá, Derciliano Lopes da Cruz, Claudio Eduardo Cavalcanti, Armando de Menezes Neto, Caroline Targino Alves da Silva, Renata Pessôa Germano Mendes, Maria Almerice Lopes da Silva, Tiago Gräf, Paola Cristina Resende, Gonzalo Bello, Michelle da Silva Barros, Wheverton Ricardo Correia do Nascimento, Rodrigo Moraes Loyo Arcoverde, Luciane Caroline Albuquerque Bezerra, Sinval Pinto Brandão Filho, Constância Flávia Junqueira Ayres, Gabriel Luz Wallau |
| EPI_ISL_510535 | Molecular Virology, Instituto Carlos Chagas / Fiocruz Paraná | Universidade Federal do Parana (UFPR) | Suzukawa,A., Tscha,M., Zanluca,C., Raboni,S., Duarte dos Santos,C. |
| EPI_ISL_513513, EPI_ISL_513515, EPI_ISL_513516, EPI_ISL_513517, EPI_ISL_513518, EPI_ISL_513519, EPI_ISL_513520, EPI_ISL_513521, EPI_ISL_513522, EPI_ISL_513523, EPI_ISL_513525, EPI_ISL_513526, EPI_ISL_513528, EPI_ISL_513530, EPI_ISL_513531, EPI_ISL_513533, EPI_ISL_513534, EPI_ISL_513535, EPI_ISL_513536, EPI_ISL_513537, EPI_ISL_513538, EPI_ISL_513540, EPI_ISL_513541, EPI_ISL_513542, EPI_ISL_513543, EPI_ISL_513544, EPI_ISL_513545, EPI_ISL_513547, EPI_ISL_513548, EPI_ISL_513549, EPI_ISL_513550, EPI_ISL_513551, EPI_ISL_513552, EPI_ISL_513553, EPI_ISL_513554, EPI_ISL_513555, EPI_ISL_513556, EPI_ISL_513558, EPI_ISL_513559, EPI_ISL_513560, EPI_ISL_513561, EPI_ISL_513562, EPI_ISL_513563, EPI_ISL_513564, EPI_ISL_513565, EPI_ISL_513566, EPI_ISL_513567, EPI_ISL_513568, EPI_ISL_513569, EPI_ISL_513570, EPI_ISL_513571, EPI_ISL_513572, EPI_ISL_513573, EPI_ISL_513574, EPI_ISL_513575, EPI_ISL_513576, EPI_ISL_513577, EPI_ISL_513579, EPI_ISL_513580, EPI_ISL_513581, EPI_ISL_513582, EPI_ISL_513583 | | | |
| see above | Programa de Oncovirologia, Instituto Nacional de Câncer | Programa de Oncovirologia, Instituto Nacional de Câncer | Juliana D. Siqueira, Livia R. Goes, Brunna M. Alves, Claudia Cicala,James Arthos, João P.B. Viola, Andreia C. de Melo, Marcelo A. Soares |
| EPI_ISL_514131 | Rondônia Central Public Health Laboratory (LACEN/RO), vinculated to State Health Secretariat of Rondônia (SESAU/RO) | Molecular Virology Laboratory of Oswaldo Cruz Foundation of Rondônia | Luan Felipo Botelho-Souza, Felipe Souza Nogueira-Lima, Tárcio Peixoto Roca, Alcione de Oliveira dos Santos, Felipe Gomes Naveca, Adriana Cristina Salvador Maia, Cicileia Correia da Silva, Aline Linhares Ferreira de Melo Mendonça, Celina Aparecida Bertoni Lugtenburg, Camila Flávia Gomes Azzi, Juliana Loca Furtado, Suelen Cavalcante, Rita de Cássia Pontello Rampazzo, Caio Henrique Nemeth Santos, Alice Paula Di Sabatino Guimarães, Jansen Fernandes de Medeiros, Fernando Rodrigues Máximo, Juan Miguel Vilallobos-Salcedo and Deusilene Souza Vieira1 |
| EPI_ISL_514132 | Rondônia Central Public Health Laboratory (LACEN/RO), vinculated to State Health Secretariat of Rondônia (SESAU/RO) | Molecular Virology Laboratory of Oswaldo Cruz Foundation of Rondônia | Luan Felipo Botelho-Souza, Felipe Souza Nogueira-Lima, Tárcio Peixoto Roca, Alcione de Oliveira dos Santos, Felipe Gomes Naveca, Adriana Cristina Salvador Maia, Cicileia Correia da Silva, Aline Linhares Ferreira de Melo Mendonça, Celina Aparecida Bertoni Lugtenburg, Camila Flávia Gomes Azzi, Juliana Loca Furtado, Suelen Cavalcante, Rita de Cássia Pontello Rampazzo, Caio Henrique Nemeth Santos, Alice Paula Di Sabatino Guimarães, Jansen Fernandes de Medeiros, Fernando Rodrigues Máximo, Juan Miguel Vilallobos-Salcedo and Deusilene Souza Vieira. |
| EPI_ISL_514133, EPI_ISL_514135, EPI_ISL_514137, EPI_ISL_514138 | Rondônia Central Public Health Laboratory (LACEN/RO), vinculated to State Health Secretariat of Rondônia (SESAU/RO) | Molecular Virology Laboratory of Oswaldo Cruz Foundation of Rondônia | Luan Felipo Botelho-Souza, Felipe Souza Nogueira-Lima, Tárcio Peixoto Roca, Alcione de Oliveira dos Santos, Felipe Gomes Naveca, Adriana Cristina Salvador Maia, Cicileia Correia da Silva, Aline Linhares Ferreira de Melo Mendonça, Celina Aparecida Bertoni Lugtenburg, Camila Flávia Gomes Azzi, Juliana Loca Furtado, Suelen Cavalcante, Rita de Cássia Pontello Rampazzo, Caio Henrique Nemeth Santos, Alice Paula Di Sabatino Guimarães, Jansen Fernandes de Medeiros, Fernando Rodrigues Máximo, Juan Miguel Vilallobos-Salcedo and Deusilene Souza Vieira |
| EPI_ISL_515521 | Hospital Municipal Dr Waldemar Tebaldi | Instituto Adolfo Lutz, Interdiciplinary Procedures Center, Strategic Laboratory | Claudio Tavares Sacchi, Claudia Regina Gonçalves, Erica Valessa Ramos Gomes |
| EPI_ISL_515522 | UPA 24HS de Itatiba | Instituto Adolfo Lutz, Interdiciplinary Procedures Center, Strategic Laboratory | Claudio Tavares Sacchi, Claudia Regina Gonçalves, Erica Valessa Ramos Gomes |
| EPI_ISL_515523 | PS Municipal Dr Lauro Ribas Braga | Instituto Adolfo Lutz, Interdiciplinary Procedures Center, Strategic Laboratory | Claudio Tavares Sacchi, Claudia Regina Gonçalves, Erica Valessa Ramos Gomes |
| EPI_ISL_515525 | National Influenza Center - Instituto Adolfo Lutz | Instituto Adolfo Lutz, Interdiciplinary Procedures Center, Strategic Laboratory | Claudio Tavares Sacchi, Claudia Regina Gonçalves, Erica Valessa Ramos Gomes |
| EPI_ISL_515527 | Hospital Santa Clara | Instituto Adolfo Lutz, Interdiciplinary Procedures Center, Strategic Laboratory | Claudio Tavares Sacchi, Claudia Regina Gonçalves, Erica Valessa Ramos Gomes |
| EPI_ISL_515528 | Hospital Sao Paulo de Ensino da Unifesp | Instituto Adolfo Lutz, Interdiciplinary Procedures Center, Strategic Laboratory | Claudio Tavares Sacchi, Claudia Regina Gonçalves, Erica Valessa Ramos Gomes |
| EPI_ISL_515543 | Serviço de Vigilância Sanitária e Epidemiológica | Instituto Adolfo Lutz, Interdiciplinary Procedures Center, Strategic Laboratory | Claudio Tavares Sacchi, Claudia Regina Gonçalves, Erica Valessa Ramos Gomes |
| EPI_ISL_515550 | UPA Vila Santa Catarina | Instituto Adolfo Lutz, Interdiciplinary Procedures Center, | Claudio Tavares Sacchi, Claudia Regina Gonçalves, Erica Valessa Ramos Gomes |

| | | | |
|---|---|---|---|
| | | Strategic Laboratory | |
| EPI_ISL_523959 | Pronto Socorro Municipal de Perus | Instituto Adolfo Lutz, Interdiciplinary Procedures Center, Strategic Laboratory | Claudio Tavares Sacchi, Claudia Regina Gonçalves, Erica Valessa Ramos Gomes |
| EPI_ISL_523961 | Pronto Socorro Municipal 21 de Junho | Instituto Adolfo Lutz, Interdiciplinary Procedures Center, Strategic Laboratory | Claudio Tavares Sacchi, Claudia Regina Gonçalves, Erica Valessa Ramos Gomes |
| EPI_ISL_523973 | PS Municipal Dona Maria Antonieta Ferreira de Barros | Instituto Adolfo Lutz, Interdiciplinary Procedures Center, Strategic Laboratory | Claudio Tavares Sacchi, Claudia Regina Gonçalves, Erica Valessa Ramos Gomes |
| EPI_ISL_523976 | Hospital Municipal do Tatuape Carmino Caricchio | Instituto Adolfo Lutz, Interdiciplinary Procedures Center, Strategic Laboratory | Claudio Tavares Sacchi, Claudia Regina Gonçalves, Erica Valessa Ramos Gomes |
| EPI_ISL_523991, EPI_ISL_523992 | Hospital Municipal Carmen Prudente | Instituto Adolfo Lutz, Interdiciplinary Procedures Center, Strategic Laboratory | Claudio Tavares Sacchi, Claudia Regina Gonçalves, Erica Valessa Ramos Gomes |
| EPI_ISL_524467 | Hospital Municipal Dr. Moysés Deutsch | Instituto Adolfo Lutz, Interdiciplinary Procedures Center, Strategic Laboratory | Claudio Tavares Sacchi, Claudia Regina Gonçalves, Erica Valessa Ramos Gomes |
| EPI_ISL_524470 | Hospital do Servidor Público Estadual Francisco Morato de Oliveira | Instituto Adolfo Lutz, Interdiciplinary Procedures Center, Strategic Laboratory | Claudio Tavares Sacchi, Claudia Regina Gonçalves, Erica Valessa Ramos Gomes |
| EPI_ISL_524784, EPI_ISL_524788, EPI_ISL_524789, EPI_ISL_524790, EPI_ISL_524791, EPI_ISL_524792, EPI_ISL_524793, EPI_ISL_524794, EPI_ISL_524795, EPI_ISL_524796, EPI_ISL_524797, EPI_ISL_524798, EPI_ISL_524799 | | | |
| see above | Evandro Chagas Institute | Evandro Chagas Institute | Santos, M.C.; Silva, A.M.; Junior, W.D.C.; Barbagelata, L.S.; Ferreira, J.A.; Sousa, E.M.A.; da Silva, P.S.; Resque, H.R; Martins, L.C.; Sousa Junior, E.C.;Viana, G.M.R |
| EPI_ISL_527858 | Pronto Atendimento Sancta Maggiore Jardim Paulista | Instituto Adolfo Lutz, Interdiciplinary Procedures Center, Strategic Laboratory | Claudio Tavares Sacchi, Claudia Regina Gonçalves, Erica Valessa Ramos Gomes |
| EPI_ISL_527864 | Hospital e Pronto Socorro Comunitário Vila Iolanda | Instituto Adolfo Lutz, Interdiciplinary Procedures Center, Strategic Laboratory | Claudio Tavares Sacchi, Claudia Regina Gonçalves, Erica Valessa Ramos Gomes |
| EPI_ISL_527867 | Pronto Socorro Municipal - Balneario São José | Instituto Adolfo Lutz, Interdiciplinary Procedures Center, Strategic Laboratory | Claudio Tavares Sacchi, Claudia Regina Gonçalves, Erica Valessa Ramos Gomes |
| EPI_ISL_527869 | Hospital Municipal Carmen Prudente | Instituto Adolfo Lutz, Interdiciplinary Procedures Center, Strategic Laboratory | Claudio Tavares Sacchi, Claudia Regina Gonçalves, Erica Valessa Ramos Gomes |
| EPI_ISL_528637, EPI_ISL_528638 | LVM/UFRJ | Bioinformatics Laboratory / LNCC | Gustavo D. P. Silva; M. Romário M. de Souza; Bruno B. Bezerra; Lucio A. Caldas; Fabio Limonte; Elena Cobos; Sharton V. A. Coelho; Luiz Almeida; Luiza Higga; Isadora A. Correa; Diana Marianni; Luciana B. Arruda; Marcelo Bozza; Orlando Ferreira; Wanderley de Souza; Ana Teresa R. Vasconcelos; Terezinha M. Castineiras; Amilcar Tanuri; Luciana J. Costa |
| EPI_ISL_534312 | Distrito Sanitario Sul | Instituto Adolfo Lutz, Interdiciplinary Procedures Center, Strategic Laboratory | Claudio Tavares Sacchi, Claudia Regina Gonçalves, Erica Valessa Ramos Gomes |
| EPI_ISL_534315 | Serviço de Verificação de Óbitos SVO Guaruulhos | Instituto Adolfo Lutz, Interdiciplinary Procedures Center, Strategic Laboratory | Claudio Tavares Sacchi, Claudia Regina Gonçalves, Erica Valessa Ramos Gomes |
| EPI_ISL_534323 | Hospital e Pronto Socorro Comunitario Vila Yolanda | Instituto Adolfo Lutz, Interdiciplinary Procedures Center, Strategic Laboratory | Claudio Tavares Sacchi, Claudia Regina Gonçalves, Erica Valessa Ramos Gomes |
| EPI_ISL_534325 | Unidade de Vigilancia em Saude de Guarulhos | Instituto Adolfo Lutz, Interdiciplinary Procedures Center, Strategic Laboratory | Claudio Tavares Sacchi, Claudia Regina Gonçalves, Erica Valessa Ramos Gomes |
| EPI_ISL_541347, EPI_ISL_541348, EPI_ISL_541349, EPI_ISL_541350, EPI_ISL_541352, EPI_ISL_541353, EPI_ISL_541356, EPI_ISL_541357, EPI_ISL_541358, EPI_ISL_541360, EPI_ISL_541361, EPI_ISL_541362, EPI_ISL_541363, EPI_ISL_541364, EPI_ISL_541365, EPI_ISL_541367, EPI_ISL_541369 | | | |
| see above | Laboratory of Respiratory Viruses and Measles, Oswaldo Cruz Institute, FIOCRUZ | Laboratory of Respiratory Viruses and Measles, Oswaldo Cruz Institute, FIOCRUZ | Paola Resende, Luciana Appolinario, Fernando Motta, Anna Carolina Paixão, Ana Carolina Mendonça, Jonathan Lopes, Marilda Siqueira |
| EPI_ISL_541370 | LACEN/SC | Laboratory of Respiratory Viruses and Measles, Oswaldo Cruz Institute, FIOCRUZ | Paola Resende, Luciana Appolinario, Fernando Motta, Anna Carolina Paixão, Ana Carolina Mendonça, Jonathan Lopes, Sandra Bianchini, Marilda Siqueira |
| EPI_ISL_541376, EPI_ISL_541382, EPI_ISL_541385, EPI_ISL_541388, EPI_ISL_541392 | LACEN/SE | Laboratory of Respiratory Viruses and Measles, Oswaldo Cruz Institute, FIOCRUZ | Paola Resende, Luciana Appolinario, Fernando Motta, Anna Carolina Paixão, Ana Carolina Mendonça, Jonathan Lopes, Clioma Santos, Marilda Siqueira |
| EPI_ISL_541397, EPI_ISL_541398, EPI_ISL_541399 | Laboratório de Virologia Comparada e Ambiental- LVCA-IOC | Laboratory of Respiratory Viruses and Measles, Oswaldo Cruz Institute, FIOCRUZ | Paola Resende, Luciana Appolinario, Tulio Machado Fumian, Tatiana Prado, Camille Ferreira Mannarino, Fernando Motta, Ana Carolina Mendonça, Marilda Siqueira, Marize Pereira Miagostovich |
| EPI_ISL_547570 | PS Municipal Dr Augusto Gomes de Mattos | Instituto Adolfo Lutz, Interdiciplinary Procedures Center, Strategic Laboratory | Claudio Tavares Sacchi, Claudia Regina Gonçalves, Erica Valessa Ramos Gomes, Karoline Rodrigues Campos |
| EPI_ISL_547574 | Hospital Universitario da USP | Instituto Adolfo Lutz, Interdiciplinary Procedures Center, Strategic Laboratory | Claudio Tavares Sacchi, Claudia Regina Gonçalves, Erica Valessa Ramos Gomes, Karoline Rodrigues Campos |
| EPI_ISL_547578 | Hospital Doutor Domingos Leonardo Cerávolo | Instituto Adolfo Lutz, Interdiciplinary Procedures Center, Strategic Laboratory | Claudio Tavares Sacchi, Claudia Regina Gonçalves, Erica Valessa Ramos Gomes, Karoline Rodrigues Campos |
| EPI_ISL_547580 | Santa Casa da Misericórdia de Presidente Prudente | Instituto Adolfo Lutz, Interdiciplinary Procedures Center, Strategic Laboratory | Claudio Tavares Sacchi, Claudia Regina Gonçalves, Erica Valessa Ramos Gomes, Karoline Rodrigues Campos |
| EPI_ISL_572364, EPI_ISL_572369, EPI_ISL_572377, EPI_ISL_572385 | LACEN/PE | WallauLab, Aggeu Magalhaes Institute | Marcelo Henrique Santos Paiva, Duschinka Ribeiro Duarte Guedes, Cássia Docena, Matheus Filgueira Bezerra, Filipe Zimmer Dezordi, Laís Ceschini Machado, Larissa Krokovsky, Elisama Helvecio, Alexandre Freitas da Silva, Luydson Richardson Silva Vasconcelos, Antonio Mauro Rezende, Severino Jefferson Ribeiro da Silva, Kamila Gaudêncio da Silva Sales, Bruna Santos Lima Figueiredo de Sá, Derciliano Lopes da Cruz, Claudio Eduardo Cavalcanti, Armando de Menezes Neto, Caroline Targino Alves da Silva, Renata Pessôa Germano Mendes, Maria Almerice Lopes da Silva, Tiago Gräf, Paola Cristina Resende, Gonzalo Bello0, Michelle da Silva Barros, Wheverton Ricardo Correia do Nascimento,, Rodrigo Moraes Loyo Arcoverde, Luciane Caroline Albuquerque Bezerra, Sinval Pinto Brandão Filho, Constância Flávia Junqueira Ayres, Gabriel Luz Wallau |
| EPI_ISL_574593, EPI_ISL_574596 | CS II Dr. Antonio Vicoso Moreira de Rezende Sumare | Instituto Adolfo Lutz, Interdiciplinary Procedures Center, Strategic Laboratory | Claudio Tavares Sacchi, Claudia Regina Gonçalves, Erica Valessa Ramos Gomes, Karoline Rodrigues Campos |
| EPI_ISL_583491 | Centro de Saude Esf IV Zona Rual Domingos de SJ Rio Pardo | Instituto Adolfo Lutz, Interdiciplinary Procedures Center, Strategic Laboratory | Claudio Tavares Sacchi, Claudia Regina Gonçalves, Erica Valessa Ramos Gomes, Karoline Rodrigues Campos |
| EPI_ISL_583493 | Vigilância em Saúde de Cajamar | Instituto Adolfo Lutz, Interdiciplinary Procedures Center, Strategic Laboratory | Claudio Tavares Sacchi, Claudia Regina Gonçalves, Erica Valessa Ramos Gomes, Karoline Rodrigues Campos |
| EPI_ISL_583495 | Serviço de Verificação de Óbitos SVO Guaruulhos | Instituto Adolfo Lutz, Interdiciplinary Procedures Center, Strategic Laboratory | Claudio Tavares Sacchi, Claudia Regina Gonçalves, Erica Valessa Ramos Gomes, Karoline Rodrigues Campos |
| EPI_ISL_603025 | UPA Central de Caraguatatuba | Instituto Adolfo Lutz, Interdiciplinary Procedures Center, Strategic Laboratory | Claudio Tavares Sacchi, Claudia Regina Gonçalves, Erica Valessa Ramos Gomes, Karoline Rodrigues Campos |

| | | | |
|---|---|---|---|
| EPI_ISL_603026 | Santa Casa da Misericórdia de Presidente Prudente | Instituto Adolfo Lutz, Interdiciplinary Procedures Center, Strategic Laboratory | Claudio Tavares Sacchi, Claudia Regina Gonçalves, Erica Valessa Ramos Gomes, Karoline Rodrigues Campos |
| EPI_ISL_603029 | Hospital Municipal Mário Gatti | Instituto Adolfo Lutz, Interdiciplinary Procedures Center, Strategic Laboratory | Claudio Tavares Sacchi, Claudia Regina Gonçalves, Erica Valessa Ramos Gomes, Karoline Rodrigues Campos |
| EPI_ISL_603031 | Santa Casa de Presidente Epitácio | Instituto Adolfo Lutz, Interdiciplinary Procedures Center, Strategic Laboratory | Claudio Tavares Sacchi, Claudia Regina Gonçalves, Erica Valessa Ramos Gomes, Karoline Rodrigues Campos |
| EPI_ISL_603032 | Santa Casa da Misericórdia de Presidente Prudente | Instituto Adolfo Lutz, Interdiciplinary Procedures Center, Strategic Laboratory | Claudio Tavares Sacchi, Claudia Regina Gonçalves, Erica Valessa Ramos Gomes, Karoline Rodrigues Campos |
| EPI_ISL_603039 | Hospital Municipal Mário Gatti | Instituto Adolfo Lutz, Interdiciplinary Procedures Center, Strategic Laboratory | Claudio Tavares Sacchi, Claudia Regina Gonçalves, Erica Valessa Ramos Gomes, Karoline Rodrigues Campos |
| EPI_ISL_613563, EPI_ISL_613564, EPI_ISL_613707, EPI_ISL_613708 | Laboratory of Molecular Biology, Blood Center of Ribeirão Preto | Laboratory of Molecular Biology, Blood Center of Ribeirão Preto, Faculty of Medicine of Ribeirão Preto, University of São Paulo | Svetoslav N Slavov, Marta Giovanetti, Vagner Fonseca, Elaine V Santos, Evandra S Rodrigues, Talita Adelino, Joilson Xavier, Glauco de Carvalho Pereira, Aparecida Y Yamamoto, Diego Villa Clé, Rodrigo T Calado; Dimas T Covas, Luiz CJ Alcantara, Simone Kashima |
| EPI_ISL_613709, EPI_ISL_613711, EPI_ISL_613951, EPI_ISL_613952, EPI_ISL_613954, EPI_ISL_613956, EPI_ISL_613961, EPI_ISL_613962 | Laboratory of Molecular Biology, Blood Center of Ribeirão Preto, Faculty of Medicine of Ribeirão Preto, University of São Paulo | Laboratory of Molecular Biology, Blood Center of Ribeirão Preto, Faculty of Medicine of Ribeirão Preto, University of São Paulo | Svetoslav N Slavov, Marta Giovanetti, Vagner Fonseca, Elaine V Santos, Evandra S Rodrigues, Talita Adelino, Joilson Xavier, Glauco de Carvalho Pereira, Aparecida Y Yamamoto, Diego Villa Clé, Rodrigo T Calado; Dimas T Covas, Luiz CJ Alcantara, Simone Kashima |
| EPI_ISL_623104, EPI_ISL_623105 | Simile Medicina Diagnóstica | Bioinformatics Laboratory / LNCC | Carolina M Voloch, Ronaldo S Francisco Jr, Luiz G P de Almeida, Otavio J. Brustolini, Cynthia C Cardoso, Alexandra L Gerber, Ana Paula de C Guimarães, Diana Mariani, Covid19-UFRJ Workgroup, Luís Cristóvão Pôrto, Renato S Aguiar, Terezinha M P P Castiñeiras, Orlando C. Ferreira, Amilcar Tanuri, Ana Tereza R de Vasconcelos |
| EPI_ISL_623106, EPI_ISL_623107, EPI_ISL_623108, EPI_ISL_623109, EPI_ISL_623110, EPI_ISL_623111, EPI_ISL_623112, EPI_ISL_623113, EPI_ISL_623114, EPI_ISL_623115, EPI_ISL_623116, EPI_ISL_623117, EPI_ISL_623118, EPI_ISL_623119, EPI_ISL_623120, EPI_ISL_623121, EPI_ISL_623124, EPI_ISL_623125, EPI_ISL_623126, EPI_ISL_623127, EPI_ISL_623128, EPI_ISL_623129, EPI_ISL_623132, EPI_ISL_623133, EPI_ISL_623134, EPI_ISL_623135, EPI_ISL_623140, EPI_ISL_623141, EPI_ISL_623142, EPI_ISL_623143, EPI_ISL_623144, EPI_ISL_623145, EPI_ISL_623146, EPI_ISL_623147, EPI_ISL_623148, EPI_ISL_623149, EPI_ISL_623150, EPI_ISL_623151, EPI_ISL_623152, EPI_ISL_623153, EPI_ISL_623154, EPI_ISL_623155, EPI_ISL_623156, EPI_ISL_623157, EPI_ISL_623158, EPI_ISL_623159, EPI_ISL_623160, EPI_ISL_623161, EPI_ISL_623162, EPI_ISL_623163, EPI_ISL_623164, EPI_ISL_623166, EPI_ISL_623167, EPI_ISL_623168, EPI_ISL_623169 | | | |
| see above | Laboratorio de Virologia Molecular / UFRJ | Bioinformatics Laboratory / LNCC | Carolina M Voloch, Ronaldo S Francisco Jr, Luiz G P de Almeida, Otavio J. Brustolini, Cynthia C Cardoso, Alexandra L Gerber, Ana Paula de C Guimarães, Diana Mariani, Covid19-UFRJ Workgroup, Luís Cristóvão Pôrto, Renato S Aguiar, Terezinha M P P Castiñeiras, Orlando C. Ferreira, Amilcar Tanuri, Ana Tereza R de Vasconcelos |
| EPI_ISL_636737, EPI_ISL_636835, EPI_ISL_636836, EPI_ISL_636837, EPI_ISL_636838 | Laboratório de Imunofarmacologia - Instituto Oswaldo Cruz | Laboratório de Imunofarmacologia - Instituto Oswaldo Cruz | Souza,T.M., Fintelman-Rodrigues,N., De Paula,A.D., Saraiva,F.B., Ferreira,M.A. and Sacramento,C.Q. |
| EPI_ISL_672664, EPI_ISL_672665, EPI_ISL_672666, EPI_ISL_672667, EPI_ISL_672668 | DB Diagnosticos do Brasil | Laboratório de Parasitologia Médica - Instituto de Medicina Tropical - Universidade de São Paulo | Brazil-UK Centre for Arbovirus Discovery Diagnosis Genomics and Epidemiology (CADDE) Genomic Network - Instituto de Medicina Tropical |
| EPI_ISL_672669 | Hospital das Clínicas da Faculdade de Medicina da Universidade de São Paulo (HC-FMUSP) | Laboratório de Parasitologia Médica - Instituto de Medicina Tropical - Universidade de São Paulo | Brazil-UK Centre for Arbovirus Discovery Diagnosis Genomics and Epidemiology (CADDE) Genomic Network - Instituto de Medicina Tropical |
| EPI_ISL_672671, EPI_ISL_672674, EPI_ISL_672675, EPI_ISL_672679, EPI_ISL_672681, EPI_ISL_672683, EPI_ISL_672684 | DB Diagnosticos do Brasil | Laboratório de Parasitologia Médica - Instituto de Medicina Tropical - Universidade de São Paulo | Brazil-UK Centre for Arbovirus Discovery Diagnosis Genomics and Epidemiology (CADDE) Genomic Network - Instituto de Medicina Tropical |
| EPI_ISL_672696 | Hospital das Clínicas da Faculdade de Medicina da Universidade de São Paulo (HC-FMUSP) | Laboratório de Parasitologia Médica - Instituto de Medicina Tropical - Universidade de São Paulo | Brazil-UK Centre for Arbovirus Discovery Diagnosis Genomics and Epidemiology (CADDE) Genomic Network - Instituto de Medicina Tropical |
| EPI_ISL_672709, EPI_ISL_672712 | Institute of Tropical Medicine at the University of São Paulo (IMT-USP) | Laboratório de Parasitologia Médica - Instituto de Medicina Tropical - Universidade de São Paulo | Brazil-UK Centre for Arbovirus Discovery Diagnosis Genomics and Epidemiology (CADDE) Genomic Network - Instituto de Medicina Tropical |
| EPI_ISL_672723, EPI_ISL_672728, EPI_ISL_672731, EPI_ISL_672736, EPI_ISL_672737, EPI_ISL_672744 | Hospital das Clínicas da Faculdade de Medicina da Universidade de São Paulo (HC-FMUSP) | Laboratório de Parasitologia Médica - Instituto de Medicina Tropical - Universidade de São Paulo | Brazil-UK Centre for Arbovirus Discovery Diagnosis Genomics and Epidemiology (CADDE) Genomic Network - Instituto de Medicina Tropical |
| EPI_ISL_672746, EPI_ISL_672751 | Institute of Tropical Medicine at the University of São Paulo (IMT-USP) | Laboratório de Parasitologia Médica - Instituto de Medicina Tropical - Universidade de São Paulo | Brazil-UK Centre for Arbovirus Discovery Diagnosis Genomics and Epidemiology (CADDE) Genomic Network - Instituto de Medicina Tropical |
| EPI_ISL_693213 | Hospital E Maternidade Municipal Governador Mario Covas | Instituto Adolfo Lutz, Interdiciplinary Procedures Center, Strategic Laboratory | Claudio Tavares Sacchi, Claudia Regina Gonçalves, Erica Valessa Ramos Gomes, Karoline Rodrigues Campos |
| EPI_ISL_693218 | Hospital Domingos Leonardo Ceravolo Presidente Prudente | Instituto Adolfo Lutz, Interdiciplinary Procedures Center, Strategic Laboratory | Claudio Tavares Sacchi, Claudia Regina Gonçalves, Erica Valessa Ramos Gomes, Karoline Rodrigues Campos |
| EPI_ISL_693219 | Santa Casa da Misericórdia de Presidente Prudente | Instituto Adolfo Lutz, Interdiciplinary Procedures Center, Strategic Laboratory | Claudio Tavares Sacchi, Claudia Regina Gonçalves, Erica Valessa Ramos Gomes, Karoline Rodrigues Campos |
| EPI_ISL_693222 | Secretaria Municipal de Saúde de Birigui | Instituto Adolfo Lutz, Interdiciplinary Procedures Center, Strategic Laboratory | Claudio Tavares Sacchi, Claudia Regina Gonçalves, Erica Valessa Ramos Gomes, Karoline Rodrigues Campos |
| EPI_ISL_693227 | UBS Vila Marchi | Instituto Adolfo Lutz, Interdiciplinary Procedures Center, Strategic Laboratory | Claudio Tavares Sacchi, Claudia Regina Gonçalves, Erica Valessa Ramos Gomes, Karoline Rodrigues Campos |
| EPI_ISL_693246 | Laboratorio Municipal de Rio Grande da Serra | Instituto Adolfo Lutz, Interdiciplinary Procedures Center, Strategic Laboratory | Claudio Tavares Sacchi, Claudia Regina Gonçalves, Erica Valessa Ramos Gomes, Karoline Rodrigues Campos |
| EPI_ISL_693247 | Secao Centro de Diagnostico Secedi | Instituto Adolfo Lutz, Interdiciplinary Procedures Center, Strategic Laboratory | Claudio Tavares Sacchi, Claudia Regina Gonçalves, Erica Valessa Ramos Gomes, Karoline Rodrigues Campos |
| EPI_ISL_693248 | Centro Municipal de Epidemiologia e Imunizações | Instituto Adolfo Lutz, Interdiciplinary Procedures Center, Strategic Laboratory | Claudio Tavares Sacchi, Claudia Regina Gonçalves, Erica Valessa Ramos Gomes, Karoline Rodrigues Campos |
| EPI_ISL_708529 | Secretária Municipal de Saude de Fernandópolis | Instituto Adolfo Lutz, Interdiciplinary Procedures Center, Strategic Laboratory | Claudio Tavares Sacchi, Claudia Regina Gonçalves, Erica Valessa Ramos Gomes, Carlos Henrique Camargo, Karoline Rodrigues Campos, Fernanda Modesto Tolentino Binhardi, Maricelia Navarro Pinheiro Flores, Marcia Maria Costa Nunes Soares, Janaina Other Martins Montanha |
| EPI_ISL_717785, EPI_ISL_717788, EPI_ISL_717789, EPI_ISL_717790 | LACEN RJ - Noel Nutels | Bioinformatics Laboratory / LNCC | Carolina M Voloch, Ronaldo da Silva F Jr, Luiz G P de Almeida, Cynthia C Cardoso, Otavio Bustrolini, Alexandra L Gerber, Ana Paula de C Guimarães, Diana Mariani, Andréa Cony Cavalcanti, Claudia dos Santos Rodrigues, Terezinha M P P Castiñeira, Amilcar Tanuri, Ana Tereza R de Vasconcelos |
| EPI_ISL_717793 | Laboratorio de Virologia Molecular / UFRJ | Bioinformatics Laboratory / LNCC | Carolina M Voloch, Ronaldo da Silva F Jr, Luiz G P de Almeida, Cynthia C Cardoso, Otavio Bustrolini, Alexandra L Gerber, Ana Paula de C Guimarães, Diana Mariani, Andréa Cony Cavalcanti, Claudia dos Santos Rodrigues, Terezinha M P P Castiñeira, Amilcar Tanuri, Ana Tereza R de Vasconcelos |
| EPI_ISL_717796, EPI_ISL_717797, EPI_ISL_717803 | LACEN RJ - Noel Nutels | Bioinformatics Laboratory / LNCC | Carolina M Voloch, Ronaldo da Silva F Jr, Luiz G P de Almeida, Cynthia C Cardoso, Otavio Bustrolini, Alexandra L Gerber, Ana Paula de C Guimarães, Diana Mariani, Andréa Cony Cavalcanti, Claudia dos Santos Rodrigues, Terezinha M P P Castiñeira, Amilcar Tanuri, Ana Tereza R de Vasconcelos |
| EPI_ISL_717832, EPI_ISL_717833, | LACEN Dr. Francisco Rimolo Neto | Bioinformatics Laboratory / LNCC | Carolina M Voloch, Ronaldo da Silva F Jr, Luiz G P de Almeida, Cynthia C Cardoso, Otavio Bustrolini, Alexandra L Gerber, Ana Paula de C Guimarães, |

| | | | |
|---|---|---|---|
| EPI_ISL_717834, EPI_ISL_717835, EPI_ISL_717836 | | | Diana Mariani, Andréa Cony Cavalcanti, Claudia dos Santos Rodrigues, Terezinha M P P Castiñeira, Amilcar Tanuri, Ana Tereza R de Vasconcelos |
| EPI_ISL_717837, EPI_ISL_717838, EPI_ISL_717839, EPI_ISL_717840 | Laboratorio de Virologia Molecular / UFRJ | Bioinformatics Laboratory / LNCC | Carolina M Voloch, Ronaldo da Silva F Jr, Luiz G P de Almeida, Cynthia C Cardoso, Otavio Bustrolini, Alexandra L Gerber, Ana Paula de C Guimarães, Diana Mariani, Andréa Cony Cavalcanti, Claudia dos Santos Rodrigues, Terezinha M P P Castiñeira, Amilcar Tanuri, Ana Tereza R de Vasconcelos |
| EPI_ISL_717841 | LACEN Dr. Francisco Rimolo Neto | Bioinformatics Laboratory / LNCC | Carolina M Voloch, Ronaldo da Silva F Jr, Luiz G P de Almeida, Cynthia C Cardoso, Otavio Bustrolini, Alexandra L Gerber, Ana Paula de C Guimarães, Diana Mariani, Andréa Cony Cavalcanti, Claudia dos Santos Rodrigues, Terezinha M P P Castiñeira, Amilcar Tanuri, Ana Tereza R de Vasconcelos |
| EPI_ISL_717842, EPI_ISL_717843, EPI_ISL_717844, EPI_ISL_717845, EPI_ISL_717846, EPI_ISL_717847, EPI_ISL_717848, EPI_ISL_717849, EPI_ISL_717850, EPI_ISL_717851, EPI_ISL_717852, EPI_ISL_717853, EPI_ISL_717854, EPI_ISL_717855, EPI_ISL_717856, EPI_ISL_717857, EPI_ISL_717858, EPI_ISL_717859, EPI_ISL_717860, EPI_ISL_717861, EPI_ISL_717862, EPI_ISL_717863, EPI_ISL_717864, EPI_ISL_717865, EPI_ISL_717866, EPI_ISL_717867, EPI_ISL_717868, EPI_ISL_717869, EPI_ISL_717870, EPI_ISL_717871, EPI_ISL_717872, EPI_ISL_717873, EPI_ISL_717874, EPI_ISL_717875, EPI_ISL_717876, EPI_ISL_717877, EPI_ISL_717880, EPI_ISL_717881, EPI_ISL_717882, EPI_ISL_717883, EPI_ISL_717884, EPI_ISL_717885, EPI_ISL_717886, EPI_ISL_717887, EPI_ISL_717888, EPI_ISL_717889, EPI_ISL_717890, EPI_ISL_717891, EPI_ISL_717892, EPI_ISL_717893, EPI_ISL_717894, EPI_ISL_717896, EPI_ISL_717897, EPI_ISL_717898 | | | |
| see above | Laboratorio de Virologia Molecular / UFRJ | Bioinformatics Laboratory / LNCC | Carolina M Voloch, Ronaldo da Silva F Jr, Luiz G P de Almeida, Cynthia C Cardoso, Otavio Bustrolini, Alexandra L Gerber, Ana Paula de C Guimarães, Diana Mariani, Andréa Cony Cavalcanti, Claudia dos Santos Rodrigues, Terezinha M P P Castiñeira, Amilcar Tanuri, Ana Tereza R de Vasconcelos |
| EPI_ISL_717900, EPI_ISL_717901, EPI_ISL_717902, EPI_ISL_717903, EPI_ISL_717904, EPI_ISL_717905, EPI_ISL_717906, EPI_ISL_717907, EPI_ISL_717908, EPI_ISL_717909 | LACEN RJ - Noel Nutels | Bioinformatics Laboratory / LNCC | Carolina M Voloch, Ronaldo da Silva F Jr, Luiz G P de Almeida, Cynthia C Cardoso, Otavio Bustrolini, Alexandra L Gerber, Ana Paula de C Guimarães, Diana Mariani, Andréa Cony Cavalcanti, Claudia dos Santos Rodrigues, Terezinha M P P Castiñeira, Amilcar Tanuri, Ana Tereza R de Vasconcelos |
| EPI_ISL_717910, EPI_ISL_717911, EPI_ISL_717912, EPI_ISL_717913, EPI_ISL_717914, EPI_ISL_717915, EPI_ISL_717916, EPI_ISL_717917, EPI_ISL_717918, EPI_ISL_717919, EPI_ISL_717958 | | | |
| see above | LACEN Dr. Francisco Rimolo Neto | Bioinformatics Laboratory / LNCC | Carolina M Voloch, Ronaldo da Silva F Jr, Luiz G P de Almeida, Cynthia C Cardoso, Otavio Bustrolini, Alexandra L Gerber, Ana Paula de C Guimarães, Diana Mariani, Andréa Cony Cavalcanti, Claudia dos Santos Rodrigues, Terezinha M P P Castiñeira, Amilcar Tanuri, Ana Tereza R de Vasconcelos |
| EPI_ISL_717959, EPI_ISL_717961 | Laboratorio de Virologia Molecular / UFRJ | Bioinformatics Laboratory / LNCC | Carolina M Voloch, Ronaldo da Silva F Jr, Luiz G P de Almeida, Cynthia C Cardoso, Otavio Bustrolini, Alexandra L Gerber, Ana Paula de C Guimarães, Diana Mariani, Andréa Cony Cavalcanti, Claudia dos Santos Rodrigues, Terezinha M P P Castiñeira, Amilcar Tanuri, Ana Tereza R de Vasconcelos |
| EPI_ISL_717962 | LACEN RJ - Noel Nutels | Bioinformatics Laboratory / LNCC | Carolina M Voloch, Ronaldo da Silva F Jr, Luiz G P de Almeida, Cynthia C Cardoso, Otavio Bustrolini, Alexandra L Gerber, Ana Paula de C Guimarães, Diana Mariani, Andréa Cony Cavalcanti, Claudia dos Santos Rodrigues, Terezinha M P P Castiñeira, Amilcar Tanuri, Ana Tereza R de Vasconcelos |
| EPI_ISL_717963, EPI_ISL_717964 | LACEN Dr. Francisco Rimolo Neto | Bioinformatics Laboratory / LNCC | Carolina M Voloch, Ronaldo da Silva F Jr, Luiz G P de Almeida, Cynthia C Cardoso, Otavio Bustrolini, Alexandra L Gerber, Ana Paula de C Guimarães, Diana Mariani, Andréa Cony Cavalcanti, Claudia dos Santos Rodrigues, Terezinha M P P Castiñeira, Amilcar Tanuri, Ana Tereza R de Vasconcelos |
| EPI_ISL_721991, EPI_ISL_721992, EPI_ISL_721997, EPI_ISL_721998, EPI_ISL_722000, EPI_ISL_722001, EPI_ISL_722008, EPI_ISL_722021, EPI_ISL_722028, EPI_ISL_722030, EPI_ISL_722041, EPI_ISL_722042, EPI_ISL_722043, EPI_ISL_722056, EPI_ISL_722059, EPI_ISL_722060, EPI_ISL_722067, EPI_ISL_722072, EPI_ISL_722081, EPI_ISL_722090, EPI_ISL_722096, EPI_ISL_722103, EPI_ISL_722106, EPI_ISL_722112, EPI_ISL_722115, EPI_ISL_722120, EPI_ISL_722126, EPI_ISL_722129 | | | |
| see above | Hospital das Clínicas Universidade de São Paulo Medical School | Laboratório de Parasitologia Médica - Instituto de Medicina Tropical - Universidade de São Paulo | Brazil-UK Centre for Arbovirus Discovery Diagnosis Genomics and Epidemiology (CADDE) Genomic Network - Instituto de Medicina Tropical |
| EPI_ISL_722130 | Instituto de Medicina Tropical Universidade de São Paulo | Laboratório de Parasitologia Médica - Instituto de Medicina Tropical - Universidade de São Paulo | Brazil-UK Centre for Arbovirus Discovery Diagnosis Genomics and Epidemiology (CADDE) Genomic Network - Instituto de Medicina Tropical |
| EPI_ISL_722136, EPI_ISL_722137, EPI_ISL_722138, EPI_ISL_722139, EPI_ISL_722140, EPI_ISL_722143, EPI_ISL_722144, EPI_ISL_722146, EPI_ISL_722147, EPI_ISL_722149, EPI_ISL_722150, EPI_ISL_722152, EPI_ISL_722153, EPI_ISL_722155, EPI_ISL_722156, EPI_ISL_722158, EPI_ISL_722160, EPI_ISL_722167, EPI_ISL_722168, EPI_ISL_722169 | | | |
| see above | DB Diagnosticos do Brasil | Laboratório de Parasitologia Médica - Instituto de Medicina Tropical - Universidade de São Paulo | Brazil-UK Centre for Arbovirus Discovery Diagnosis Genomics and Epidemiology (CADDE) Genomic Network - Instituto de Medicina Tropical |
| EPI_ISL_729794, EPI_ISL_729795, EPI_ISL_729796, EPI_ISL_729797, EPI_ISL_729798, EPI_ISL_729800, EPI_ISL_729802, EPI_ISL_729807, EPI_ISL_729809, EPI_ISL_729810, EPI_ISL_729812, EPI_ISL_729814, EPI_ISL_729816, EPI_ISL_729817, EPI_ISL_729818, EPI_ISL_729819, EPI_ISL_729820, EPI_ISL_729821, EPI_ISL_729823, EPI_ISL_729825, EPI_ISL_729826, EPI_ISL_729827, EPI_ISL_729828, EPI_ISL_729829, EPI_ISL_729830, EPI_ISL_729831, EPI_ISL_729832, EPI_ISL_729833, EPI_ISL_729837, EPI_ISL_729838, EPI_ISL_729839, EPI_ISL_729841, EPI_ISL_729842, EPI_ISL_729843, EPI_ISL_729844, EPI_ISL_729849, EPI_ISL_729851, EPI_ISL_729857, EPI_ISL_729858, EPI_ISL_729860 | | | |
| see above | Laboratorio Central de Saude Publica do Estado do Rio Grande do Sul (LACEN-RS) | Laboratory of Respiratory Viruses and Measles, Oswaldo Cruz Institute, FIOCRUZ | Paola Resende, Luciana Appolinario, Fernando Motta, Anna Carolina Paixão, Ana Carolina Mendonça, Tatiana Schaffer Gregianini, Marilda Tereza Mar da Rosa, Marilda Siqueira |
| EPI_ISL_735407 | Santa Casa de Marilia | Instituto Adolfo Lutz, Interdiciplinary Procedures Center, Strategic Laboratory | Claudio Tavares Sacchi, Claudia Regina Gonçalves, Erica Valessa Ramos Gomes, Karoline Rodrigues Campos |
| EPI_ISL_735410 | Instituto Adolfo Lutz - Regional de Rio Claro | Instituto Adolfo Lutz, Interdiciplinary Procedures Center, Strategic Laboratory | Claudio Tavares Sacchi, Claudia Regina Gonçalves, Erica Valessa Ramos Gomes, Karoline Rodrigues Campos |
| EPI_ISL_735414, EPI_ISL_735415 | Unidade de Pronto Atendimento de Agenor de Campos | Instituto Adolfo Lutz, Interdiciplinary Procedures Center, Strategic Laboratory | Claudio Tavares Sacchi, Claudia Regina Gonçalves, Erica Valessa Ramos Gomes, Karoline Rodrigues Campos |
| EPI_ISL_735416 | Centro de Saude II Dr Jose Paione Mococa | Instituto Adolfo Lutz, Interdiciplinary Procedures Center, Strategic Laboratory | Claudio Tavares Sacchi, Claudia Regina Gonçalves, Erica Valessa Ramos Gomes, Karoline Rodrigues Campos |
| EPI_ISL_735425 | Hospital e Maternidade Sao Lucas | Instituto Adolfo Lutz, Interdiciplinary Procedures Center, Strategic Laboratory | Claudio Tavares Sacchi, Claudia Regina Gonçalves, Erica Valessa Ramos Gomes, Karoline Rodrigues Campos |
| EPI_ISL_735427, EPI_ISL_735430 | Instituto Adolfo Lutz - Regional de Santos | Instituto Adolfo Lutz, Interdiciplinary Procedures Center, Strategic Laboratory | Claudio Tavares Sacchi, Claudia Regina Gonçalves, Erica Valessa Ramos Gomes, Karoline Rodrigues Campos |
| EPI_ISL_755649 | Instituto Adolfo Lutz - Regional de Santo Andre | Instituto Adolfo Lutz, Interdiciplinary Procedures Center, Strategic Laboratory | Claudio Tavares Sacchi, Claudia Regina Gonçalves, Erica Valessa Ramos Gomes, Karoline Rodrigues Campos |
| EPI_ISL_755652 | Lab LOC - Itapecerica da Serra | Instituto Adolfo Lutz, Interdiciplinary Procedures Center, Strategic Laboratory | Claudio Tavares Sacchi, Claudia Regina Gonçalves, Erica Valessa Ramos Gomes, Karoline Rodrigues Campos |
| EPI_ISL_756293 | Center for Biotechnology and Cell Therapy, São Rafael Hospital, Salvador, Brazil | Center for Biotechnology and Cell Therapy, São Rafael Hospital, Salvador, Brazil | Carolina Kymie Vasques Nonaka, Marília Miranda Franco, Tiago Gräf, Ana Verena Almeida Mendes, Renato Santana de Aguiar, Marta Giovanetti, Bruno Solano de Freitas Souza |
| EPI_ISL_770551, EPI_ISL_770574, EPI_ISL_770575 | Laboratório de Microbiologia Molecular - Universidade FEEVALE | Bioinformatics Laboratory / LNCC | Felipe Benites, Fernando Rosado Spilki, Alana Witt Hansen, Juliane Deise Fleck, Juliana Schons, Meriane Demoliner, Ana Karolina Eisen Antunes, Fagner Henrique Heldt, Larissa Mallmann, Bruna Hermann, Ana Luiza Ziulkoski, Vyctoria Goes, Karoline Schallenberger, Matheus Nunes Weber, Paula Rodrigues de Almeida, Alessandra Pavan Lamarca da Silva, Ronaldo da Silva F Jr , Luiz G P de Almeida, Alexandra L Gerber , Ana Paula de C Guimarães,Ana Tereza R de Vasconcelos |
| EPI_ISL_776751, EPI_ISL_776754, EPI_ISL_776759, EPI_ISL_776762 | Instituto Adolfo Lutz - Central | Instituto Adolfo Lutz, Interdiciplinary Procedures Center, Strategic Laboratory | Claudio Tavares Sacchi, Claudia Regina Gonçalves, Erica Valessa Ramos Gomes, Karoline Rodrigues Campos |
| EPI_ISL_779157, EPI_ISL_779158, EPI_ISL_779164 | Laboratório de Microbiologia Molecular - Universidade FEEVALE | Bioinformatics Laboratory / LNCC | Felipe Benites, Fernando Rosado Spilki, Alana Witt Hansen, Juliane Deise Fleck, Juliana Schons, Meriane Demoliner, Ana Karolina Eisen Antunes, Fagner Henrique Heldt, Larissa Mallmann, Bruna Hermann, Ana Luiza Ziulkoski, Vyctoria Goes, Karoline Schallenberger, Matheus Nunes Weber, Paula Rodrigues de Almeida, Alessandra Pavan Lamarca da Silva, Ronaldo da Silva F Jr , Luiz G P de Almeida, Alexandra L Gerber , Ana Paula de C Guimarães,Ana Tereza R de Vasconcelos |
| EPI_ISL_792105 | Instituto Adolfo Lutz - Central | Instituto Adolfo Lutz, Interdiciplinary Procedures Center, Strategic Laboratory | Claudio Tavares Sacchi, Claudia Regina Gonçalves, Erica Valessa Ramos Gomes, Karoline Rodrigues Campos |
| EPI_ISL_792561, EPI_ISL_792571, EPI_ISL_792572, EPI_ISL_792573, EPI_ISL_792579, EPI_ISL_792583, EPI_ISL_792588, EPI_ISL_792589, EPI_ISL_792593, EPI_ISL_792594, EPI_ISL_792596, EPI_ISL_792600, EPI_ISL_792601, EPI_ISL_792602, EPI_ISL_792603, EPI_ISL_792608, EPI_ISL_792610, EPI_ISL_792612, EPI_ISL_792623, EPI_ISL_792637 | | | |

| | | | |
|---|---|---|---|
| see above | LACEN-PB | Laboratory of Respiratory Viruses and Measles, Oswaldo Cruz Institute, FIOCRUZ | Paola Resende, Luciana Appolinario, Fernando Motta, Anna Carolina Paixao, Ana Carolina Mendonca, João Felipe Bezerra, Romero Henrique Teixeira de Vasconcelos, Dalane Loudal Florentino Teixeira, Thiago Franco de Oliveira Carneiro, Marilda Siqueira |
| EPI_ISL_792640, EPI_ISL_792644 | LACEN-AL | Laboratory of Respiratory Viruses and Measles, Oswaldo Cruz Institute, FIOCRUZ | Paola Resende, Luciana Appolinario, Fernando Motta, Anna Carolina Paixao, Ana Carolina Mendonca, Anderson Brandao Leite, Marilda Siqueira |
| EPI_ISL_792648 | LACEN-PR | Laboratory of Respiratory Viruses and Measles, Oswaldo Cruz Institute, FIOCRUZ | Paola Resende, Luciana Appolinario, Fernando Motta, Anna Carolina Paixao, Ana Carolina Mendonca, Maria do Carmo Debur, Irina Nastassja Riediger, Marilda Siqueira |
| EPI_ISL_804822, EPI_ISL_804828, EPI_ISL_804831, EPI_ISL_804838 | DB Diagnosticos do Brasil | Laboratório de Parasitologia Médica - Instituto de Medicina Tropical - Universidade de São Paulo | Nuno Faria, Ingra Morales Claro, Darlan Candido, Lucas A. Moyses Franco, Pamela dos Santos Andrade, Thais de Moura Coletti, Camila A. Maia da Silva, Flavia Cristina Sales, Erika Regina Manuli, Renato A. Santana, Nelson Gaburo, Cecília da Cunha Camilo, Nelson Abrahim Fraiji, Myuki Alfaia Esashika Crispim, Maria do Perpétuo Socorro Sampaio Carvalho, Andrew Rambaut, Nick Loman, Oliver G. Pybus, Ester C. Sabino; DB; HEMOAM; CDL; CADDE Genomic Network. |
| EPI_ISL_831474, EPI_ISL_831646, EPI_ISL_831678, EPI_ISL_831683, EPI_ISL_831685, EPI_ISL_831892, EPI_ISL_831898, EPI_ISL_831940, EPI_ISL_832012 | Laboratório de Microbiologia Molecular - Universidade FEEVALE | Universidade Federal de Ciências da Saúde de Porto Alegre | Vinícius Bonetti Franceschi, Amanda de Menezes Mayer, Gabriel Dickin Caldana, Carla Andretta Moreira Neves, Patrícia Aline Gröhs Ferrareze, Gabriela Bettella Cybis, Ricardo Ariel Zimerman, Lívia Kmetzsch, Fernando Rosado Spilki, Claudia Elizabeth Thompson |
| EPI_ISL_833135 | Laboratorio de Ecologia de Doencas Transmissiveis na Amazonia, Instituto Leonidas e Maria Deane - Fiocruz Amazonia | Laboratorio de Ecologia de Doencas Transmissiveis na Amazonia, Instituto Leonidas e Maria Deane - Fiocruz Amazonia | Valdinete Nascimento, Victor Souza, André Corado, Fernanda Nascimento, George Silva, Ágatha Costa, Debora Duarte, Karina Pessoa, Matilde Mejía, Luciana Gonçalves, Maria Júlia Brandão, Michele Jesus, Felipe Naveca |
| EPI_ISL_833155, EPI_ISL_833159 | Instituto Adolfo Lutz - Central | Instituto Adolfo Lutz, Interdiciplinary Procedures Center, Strategic Laboratory | Claudio Tavares Sacchi, Claudia Regina Gonçalves, Erica Valessa Ramos Gomes, Karoline Rodrigues Campos |
| EPI_ISL_833163 | Instituto Adolfo Lutz - Regional de Marilia | Instituto Adolfo Lutz, Interdiciplinary Procedures Center, Strategic Laboratory | Claudio Tavares Sacchi, Claudia Regina Gonçalves, Erica Valessa Ramos Gomes, Karoline Rodrigues Campos |
| EPI_ISL_848556, EPI_ISL_848561, EPI_ISL_848564, EPI_ISL_848567, EPI_ISL_848568, EPI_ISL_848569, EPI_ISL_848570, EPI_ISL_848572, EPI_ISL_848573, EPI_ISL_848574, EPI_ISL_848575, EPI_ISL_848576, EPI_ISL_848577, EPI_ISL_848578, EPI_ISL_848579, EPI_ISL_848580, EPI_ISL_848581, EPI_ISL_848584, EPI_ISL_848591, EPI_ISL_848598, EPI_ISL_848601, EPI_ISL_848603, EPI_ISL_848609, EPI_ISL_848610, EPI_ISL_848612, EPI_ISL_848613, EPI_ISL_848614, EPI_ISL_848616, EPI_ISL_848626, EPI_ISL_848627 | | | |
| see above | Evandro Chagas Institute | Evandro Chagas Institute | Santos, M.C.; Silva, A.M.; Junior, W.D.C.; Barbagelata, L.S.; Ferreira, J.A.; Sousa, E.M.A.; da Silva, P.S.; Pinheiro, K.C.; L.C.; Sousa Junior, E.C. |
| EPI_ISL_861635 | Hospital e Maternidade Madre Theodora | Instituto Adolfo Lutz, Interdiciplinary Procedures Center, Strategic Laboratory | Claudio Tavares Sacchi, Claudia Regina Gonçalves, Erica Valessa Ramos Gomes, Karoline Rodrigues Campos |
| EPI_ISL_861638 | Hospital Municipal Dr. Moyses Deutsch | Instituto Adolfo Lutz, Interdiciplinary Procedures Center, Strategic Laboratory | Claudio Tavares Sacchi, Claudia Regina Gonçalves, Erica Valessa Ramos Gomes, Karoline Rodrigues Campos |
| EPI_ISL_861642 | Instituto Adolfo Lutz - Central | Instituto Adolfo Lutz, Interdiciplinary Procedures Center, Strategic Laboratory | Claudio Tavares Sacchi, Claudia Regina Gonçalves, Erica Valessa Ramos Gomes, Karoline Rodrigues Campos |
| EPI_ISL_861651 | AMA Jardim Brasil | Instituto Adolfo Lutz, Interdiciplinary Procedures Center, Strategic Laboratory | Claudio Tavares Sacchi, Claudia Regina Gonçalves, Erica Valessa Ramos Gomes, Karoline Rodrigues Campos |
| EPI_ISL_861653 | Hospital Santa Virgínia | Instituto Adolfo Lutz, Interdiciplinary Procedures Center, Strategic Laboratory | Claudio Tavares Sacchi, Claudia Regina Gonçalves, Erica Valessa Ramos Gomes, Karoline Rodrigues Campos |
| EPI_ISL_861662 | CS I Tacito Leite de Carvalho e Silva | Instituto Adolfo Lutz, Interdiciplinary Procedures Center, Strategic Laboratory | Claudio Tavares Sacchi, Claudia Regina Gonçalves, Erica Valessa Ramos Gomes, Karoline Rodrigues Campos |
| EPI_ISL_861664 | Instituto Adolfo Lutz - Regional de Campinas | Instituto Adolfo Lutz, Interdiciplinary Procedures Center, Strategic Laboratory | Claudio Tavares Sacchi, Claudia Regina Gonçalves, Erica Valessa Ramos Gomes, Karoline Rodrigues Campos |
| EPI_ISL_861670 | Instituto Adolfo Lutz - Regional de Taubate | Instituto Adolfo Lutz, Interdiciplinary Procedures Center, Strategic Laboratory | Claudio Tavares Sacchi, Claudia Regina Gonçalves, Erica Valessa Ramos Gomes, Karoline Rodrigues Campos |
| EPI_ISL_861678 | Laboratorio Municipal de Guarulhos | Instituto Adolfo Lutz, Interdiciplinary Procedures Center, Strategic Laboratory | Claudio Tavares Sacchi, Claudia Regina Gonçalves, Erica Valessa Ramos Gomes, Karoline Rodrigues Campos |
| EPI_ISL_861869, EPI_ISL_861871, EPI_ISL_861884, EPI_ISL_861891, EPI_ISL_861893, EPI_ISL_861897, EPI_ISL_861898, EPI_ISL_861899, EPI_ISL_861907, EPI_ISL_861908, EPI_ISL_861910, EPI_ISL_861915 | | | |
| see above | LATE - Laboratório de Técnicas Especiais - Hospital Israelita Albert Einstein | LATE - Laboratório de Técnicas Especiais - Hospital Israelita Albert Einstein | Deyvid Amgarten, Fernanda de Mello Malta, Raquel Riyuzo, Ana Paula Moreira Salles, Pedro Henrique Sebe Rodrigues, João Renato Rebello Pinho |
| EPI_ISL_904021, EPI_ISL_904022, EPI_ISL_904026, EPI_ISL_904029, EPI_ISL_904032, EPI_ISL_904034 | DB Diagnosticos do Brasil | Laboratório de Parasitologia Médica - Instituto de Medicina Tropical - Universidade de São Paulo | Brazil-UK Centre for Arbovirus Discovery Diagnosis Genomics and Epidemiology (CADDE) Genomic Network - Instituto de Medicina Tropical |
| EPI_ISL_918513 | LACEN - Laboratório Central de Saúde Pública do Roraima | Evandro Chagas Institute | Santos, M.C.; Silva, A.M.; Junior, W.D.C.; Barbagelata, L.S.; Ferreira, J.A.; Sousa, E.M.A.; da Silva, P.S.; Pinheiro, K.C.; L.C.; Sousa Junior, E.C. |
| EPI_ISL_918524, EPI_ISL_918525 | LACEN - Laboratório Central de Saúde Pública do Para | Evandro Chagas Institute | Santos, M.C.; Silva, A.M.; Junior, W.D.C.; Barbagelata, L.S.; Ferreira, J.A.; Sousa, E.M.A.; da Silva, P.S.; Pinheiro, K.C.; L.C.; Sousa Junior, E.C. |
| EPI_ISL_918535 | LACEN - Laboratório Central de Saúde Pública do Amazonas | Evandro Chagas Institute | Santos, M.C.; Silva, A.M.; Junior, W.D.C.; Barbagelata, L.S.; Ferreira, J.A.; Sousa, E.M.A.; da Silva, P.S.; Pinheiro, K.C.; L.C.; Sousa Junior, E.C. |
| EPI_ISL_925846 | LACEN - Laboratório Central de Saúde Pública do Amazonas | Evandro Chagas Institute Virology | Santos, M.C.; Silva, A.M.; Junior, W.D.C.; Barbagelata, L.S.; Ferreira, J.A.; Sousa, E.M.A.; da Silva, P.S.; Pinheiro, K.C.; L.C.; Sousa Junior, E.C. |
| EPI_ISL_930854, EPI_ISL_930855, EPI_ISL_930858 | Central Laboratory of Public Health of Rio Grande do Sul (Lacen-RS) | State Center for Health Surveillance of the Health Department of the State of Rio Grande do Sul (CEVS/SES-RS) | Barcellos R, Campos A, Dornelles C, Godinho F, Gonzalez A, Gregianini T, Molina C, Salvato R, Schaurich A, |
| EPI_ISL_940613 | LACEN-PI DR. Costa Alvarenga | Instituto Adolfo Lutz, Interdiciplinary Procedures Center, Strategic Laboratory | Claudio Tavares Sacchi, Claudia Regina Gonçalves, Erica Valessa Ramos Gomes, Karoline Rodrigues Campos |
| EPI_ISL_940629 | Hospital Municipal Josanias Castanha Braga | Instituto Adolfo Lutz, Interdiciplinary Procedures Center, Strategic Laboratory | Claudio Tavares Sacchi, Claudia Regina Gonçalves, Erica Valessa Ramos Gomes, Karoline Rodrigues Campos |
| EPI_ISL_942374, EPI_ISL_942375, EPI_ISL_942407, EPI_ISL_942897, EPI_ISL_942930, EPI_ISL_942931 | Central Laboratory of Public Health of Rio Grande do Sul (Lacen-RS) | State Center for Health Surveillance of the Health Department of the State of Rio Grande do Sul (CEVS/SES-RS) | Barcellos R, Campos A, Crescente L, Da Silva A, Dornelles C, Fonseca V, Garay L, Godinho F, Gonzalez A, Gregianini T, Molina C, Salvato R, Schaurich A |
| EPI_ISL_943574, EPI_ISL_943575, EPI_ISL_943576, EPI_ISL_943577, EPI_ISL_943579, EPI_ISL_943582, EPI_ISL_943583, EPI_ISL_943585, EPI_ISL_943590, EPI_ISL_943591, EPI_ISL_943592, EPI_ISL_943593, EPI_ISL_943594, EPI_ISL_943598, EPI_ISL_943601, EPI_ISL_943605, EPI_ISL_943613 | | | |
| see above | Central Laboratory of Public Health of Rio Grande do Sul (Lacen-RS) | State Center for Health Surveillance of the Health Department of the State of Rio Grande do Sul (CEVS/SES-RS) | Aline Campos, Amanda da Silva, Anelise Schaurich, Claudia Dornelles, Cynthia Molina, Fernanda Godinho, Lara Crescente, Leticia Garay, Regina Barcellos, Richard Salvato, Tatiana Gregianini, Vagner Fonseca |
| EPI_ISL_943980, EPI_ISL_943982 | LACEN do Estado de Tocantins | Instituto Adolfo Lutz, Interdiciplinary Procedures Center, Strategic Laboratory | Claudio Tavares Sacchi, Claudia Regina Gonçalves, Erica Valessa Ramos Gomes, Karoline Rodrigues Campos |
| EPI_ISL_977479 | Lab Loc - Itapecerica da Serra | Instituto Adolfo Lutz, Interdiciplinary Procedures Center, Strategic Laboratory | Claudio Tavares Sacchi, Claudia Regina Gonçalves, Erica Valessa Ramos Gomes, Karoline Rodrigues Campos |
| EPI_ISL_977482 | Instituto Adolfo Lutz - Regional de Aracatuba | Instituto Adolfo Lutz, Interdiciplinary Procedures Center, Strategic Laboratory | Claudio Tavares Sacchi, Claudia Regina Gonçalves, Erica Valessa Ramos Gomes, Karoline Rodrigues Campos |

| | | | |
|---|---|---|---|
| EPI_ISL_977486 | Instituto Adolfo Lutz - Regional de Santo Andre | Instituto Adolfo Lutz, Interdiciplinary Procedures Center, Strategic Laboratory | Claudio Tavares Sacchi, Claudia Regina Gonçalves, Erica Valessa Ramos Gomes, Karoline Rodrigues Campos |
| EPI_ISL_977490 | Hospital Municipal Guido Guida | Instituto Adolfo Lutz, Interdiciplinary Procedures Center, Strategic Laboratory | Claudio Tavares Sacchi, Claudia Regina Gonçalves, Erica Valessa Ramos Gomes, Karoline Rodrigues Campos |
| EPI_ISL_978488, EPI_ISL_978489, EPI_ISL_978491, EPI_ISL_978492, EPI_ISL_978494, EPI_ISL_978495, EPI_ISL_978497, EPI_ISL_978499, EPI_ISL_978502, EPI_ISL_978503, EPI_ISL_978505, EPI_ISL_978507, EPI_ISL_978508, EPI_ISL_978510, EPI_ISL_978513, EPI_ISL_978514, EPI_ISL_978516, EPI_ISL_978522, EPI_ISL_978526, EPI_ISL_978528, EPI_ISL_978530, EPI_ISL_978531, EPI_ISL_978532 | | | |
| see above | Central Public Health Laboratory - LACEN -Bahia, Salvador, Brazil | Central Public Health Laboratory - LACEN -Bahia, Salvador, Brazil | Stephane Tosta, Luciana Oliveira, Vanessa Nardy,Patrícia Cajado,Marcela Gómez, Breno Dominguez, Jaqueline Gomes, Vagner Fonseca,Marta Giovanetti,Luiz Alcantara, Felicidade Pereira, Arabela Leal |
| EPI_ISL_984251, EPI_ISL_984256 | IAL Regional de Marilia | Instituto Adolfo Lutz, Interdiciplinary Procedures Center, Strategic Laboratory | Claudio Tavares Sacchi, Claudia Regina Gonçalves, Erica Valessa Ramos Gomes, Karoline Rodrigues Campos |
| EPI_ISL_985174 | Instituto Adolfo Lutz - Regional de Taubate | Instituto Adolfo Lutz, Interdiciplinary Procedures Center, Strategic Laboratory | Claudio Tavares Sacchi, Claudia Regina Gonçalves, Erica Valessa Ramos Gomes, Karoline Rodrigues Campos |
| EPI_ISL_985176 | Instituto Adolfo Lutz Central | Instituto Adolfo Lutz, Interdiciplinary Procedures Center, Strategic Laboratory | Claudio Tavares Sacchi, Claudia Regina Gonçalves, Erica Valessa Ramos Gomes, Karoline Rodrigues Campos |
